# Kinetic and Thermodynamic Insights on Agonist Interactions with the Parathyroid Hormone Receptor-1 from a New NanoBRET assay

**DOI:** 10.1101/2022.07.25.500850

**Authors:** Zhen Yu, Brian P. Cary, Tae Wook Kim, Kevin D. Nguyen, Thomas J. Gardella, Samuel H. Gellman

## Abstract

Polypeptides that activate the parathyroid hormone receptor-1 (PTHR1) are important in human physiology and medicine. Most previous studies of peptide binding to this receptor have involved displacement of a radiolabeled ligand. We report a new assay format based on bioluminescence resonance energy transfer (BRET). Fusion of a nanoluciferase (nLuc) unit to the N-terminus of the PTHR1 allows direct detection of binding by an agonist peptide bearing a tetramethylrhodamine (TMR) unit. Affinity measurements from the BRET assay align well with results previously obtained via radioligand displacement. The BRET assay offers substantial operational benefits relative to affinity measurements involving radioactive compounds. The convenience of the new assay allowed us to explore several questions raised by earlier reports. For example, we show that although the first two residues of PTH(1-34) (the drug teriparatide) are critical for PTHR1 activation, these two residues contribute little or nothing to affinity. Comparisons among the well-studied agonists PTH(1-34), PTHrP(1-34) and “long-acting PTH” (LA-PTH) reveal that the high affinity of LA-PTH arises largely from a diminished rate constant for dissociation relative to the other two.

## INTRODUCTION

The parathyroid hormone receptor-1 (PTHR1) is a family B G protein-coupled receptor (GPCR) that is targeted by several polypeptide drugs.^1–3^ Activation of the PTHR1 helps to regulate blood concentrations of calcium and phosphate ions, and this receptor plays important roles in bone physiology. The PTHR1 has two natural agonists, parathyroid hormone (PTH) and parathyroid hormone-related protein (PTHrP). PTH(1-34), a synthetic N-terminal fragment of PTH that matches the activity of the full-length hormone, is used to treat osteoporosis.^4^ Abaloparatide (ABL), a synthetic analogue of PTHrP(1-34), is used clinically for the same purpose.^5^ Full-length recombinant PTH is used to treat hypoparathyroidism.^6^ Understanding how agonists bind to and activate the PTHR1 represents a fundamental challenge in terms of biological signal transduction; such understanding could provide a basis for future therapeutic development. Here we describe a new type of assay for characterizing agonist-PTHR1 interactions, and we show how this new tool can be used to address fundamental questions regarding activation of this receptor.

As with other GPCRs, agonist binding to the PTHR1 alters the conformational equilibrium of the receptor.^7,8^ This structural change is registered by cytoplasmic proteins that engage the inward-facing surface of the receptor, including G proteins, β-arrestins and GPCR kinases (GRKs). Family B GPCRs feature a large extracellular domain (ECD) that engages the C-terminal portion of the long polypeptide agonist.^8–12^ The N-terminal region of the agonist binds into the core of the family B GPCR transmembrane domain (TMD), which contains the seven-helix bundle tertiary structure that is characteristic of all GPCRs.^13,14^

Strategies to monitor GPCR activation are valuable for characterizing relationships between agonist structure and function. Many cell-based assay formats for GPCR activation feature a convenient optical readout. Agonists of the PTHR1 and other receptors that activate G_S_, for example, are often characterized by their ability to stimulate production of the intracellular second messenger cAMP, which can be detected via luminescence generated by cAMP-binding proteins such as GloSensor.^15^ Direct recruitment of a partner protein (G protein, β-arrestin, GRK) to the intracellular surface of a GPCR can be detected via bioluminescence resonance energy transfer (BRET) if the receptor and partner are fused to complementary BRET partners, such as a luciferase donor and a fluorescent protein acceptor.^16–18^ Other optical readouts, such as fluorescence resonance energy transfer (FRET), have been harnessed for this purpose as well.^19,20^

Relative affinities among agonists for a given receptor need not correlate with relative agonist potencies; therefore, it is valuable to measure agonist binding directly, as a complement to activity measurements. Agonist binding to cognate GPCRs is commonly characterized with radioactive ligands, either through direct binding or competition with a radiolabeled tracer.^21–27^ Handling radioactive materials is cumbersome, however, which has prompted a search for more convenient binding assays. Agonist binding to a few GPCRs has recently been characterized via BRET by using a receptor construct bearing a nanoluciferase (nLuc) domain at the N-terminus in conjunction with ligands covalently attached to a complementary fluorophore, such as tetramethylrhodamine (TMR).^28,29^ This approach does not require radiolabeled compounds and involves less sample manipulation relative to radioligand displacement assays. In addition, the BRET assay format allows real-time measurements. Here we describe application this strategy to the PTHR1 (Figure 1A).

**Figure 1.**
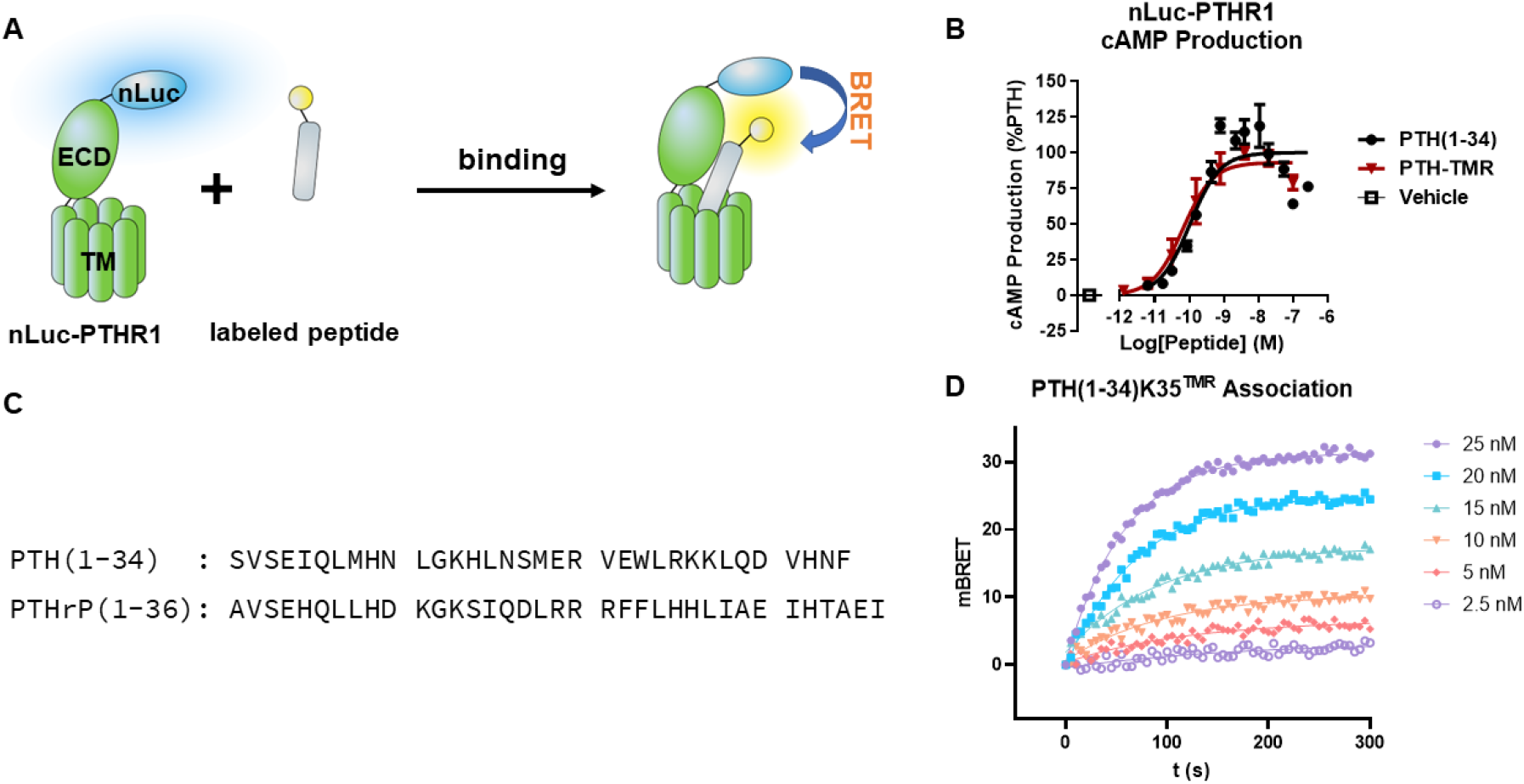
(A) Cartoon depiction of the NanoBRET binding assay. Nanoluciferase (nLuc) is attached to the extracellular side of the PTHR1, which allows energy transfer to a BRET acceptor that is covalently linked to the bound peptide. (B) cAMP production by nLuc-PTHR1-expressing cells activated by either PTH(1-34) or PTH(1-34)K35^TMR^. (C) Sequences of PTHR1 agonists. All peptides used in this work were prepared as C-terminal amides. (D) Association curves for PTH(1-34)K35^TMR^ binding to the nLuc-PTHR1 at different peptide concentrations measured over 5 min.

## RESULTS AND DISCUSSION

### An nLuc-PTHR1 fusion protein enables measurement of binding kinetics and affinities, providing results that are consistent with previous reports involving other assay formats

DNA coding for the human PTHR1 with nLuc fused to the N-terminus was transiently expressed in GS22 cells. These HEK293-derived cells stably express the GloSensor F-22 construct (Promega), which allows detection of intracellular production of cAMP. PTH(1-34) caused potent stimulation of cAMP production (EC_50_ = 0.1 nM) in these cells (Figure 1B; sequence in Figure 1C), and this activity was comparable to the activity observed for PTH(1-34) with cells transiently expressing the native human PTHR1 (EC_50_ = 0.3 nM). This agonist peptide and all others discussed below were synthesized as C-terminal amides. This comparison indicates that the N-terminal nLuc domain does not interfere with receptor activation, at least as measured by cAMP production.

To monitor agonist-receptor association, we treated cells expressing the nLuc-PTHR1 with NaN_3_ for 30 min before introducing TMR-labeled agonist and luciferase substrate (h-coelenterazine). Azide pretreatment was used to halt cellular metabolism and prevent the internalization of receptor-agonist complexes, which normally occurs shortly after agonist engagement. The lack of internalization should ensure that ligand-receptor interaction occurs solely on the cell surface, and that the ligand concentration and the total receptor density remain largely unaffected for the duration of the experiment. Ligand-receptor binding led to a rapid rise in the BRET signal, which reached a plateau within five minutes (Figure 1D). The plateau level varied with agonist concentration. These data were fitted to a pseudo-first order binding model (Eqs 1 and 2; peptide in vast excess) to generate rate constants *k*_*on*_ and *k*_*off*_. The dissociation constant, K_D_, was calculated from these rate constants, *k*_*off*_/*k*_*on*_ (Table 1).

**Table 1.** Kinetic constants and dissociation constants of various PTHR1 binding ligands as measured by either direct binding or competition binding.

| Direct Binding | $k_{on} ( \times 10^5 \text{ s}^{-1}\text{M}^{-1} )$ | $k_{off} ( \text{s}^{-1} )$ | $K_D ( \text{nM} )$ |
| --- | --- | --- | --- |
| PTH(1-34)-35K<sup>TMR</sup> | 4.4 ± 0.8 | 0.0051 ± 0.0006 | 12 ± 2 |
| PTH(1-34)-N33K<sup>TMR</sup> | 4.0 ± 0.6 | 0.0043 ± 0.002 | 11 ± 4 |
| PTHrP(1-34)-35K<sup>TMR</sup> | 2.1 ± 0.6 | 0.010 ± 0.003 | 48 ± 20 |
| PTHrP(1-34)-13K<sup>TMR</sup> | 2.4 ± 0.7 | 0.011 ± 0.001 | 53 ± 20 |
| LA-PTH(1-34)-35K<sup>TMR</sup> | 2.3 ± 0.4 | 0.00056 ± 0.0002 | 2.4 ± 0.8 |
| M-PTH(1-14)-15K<sup>TMR</sup> | 0.77 ± 0.2 | 0.024 ± 0.001 | 310 ± 80 |
| **Competition** |  |  |  |
| PTH(1-34) | 2.5 ± 1 | 0.0038 ± 0.001 | 16 ± 2 |
| PTHrP(1-34) | 2.4 ± 0.6 | 0.024 ± 0.004 | 73 ± 40 |
| **nLuc-ΔECD-PTHR1** |  |  |  |
| LA-PTH(1-34)-35K<sup>TMR</sup> | 1.3 ± 0.1 | 0.0097 ± 0.003 | 72 ± 20 |
| M-PTH(1-14)-15K<sup>TMR</sup> | 1.8 ± 0.5 | 0.018 ± 0.003 | 106 ± 20 |

The kinetics and affinity parameters of various ligands measured by different experimental approaches. *Direct binding* Measurements on nLuc-PTHR1 were obtained by treating HEK293 cells stably expressing nLuc-PTHR1 with TMR-labeled ligands in the presence of the substrate, coelenterazine-h, and monitoring the BRET response. *Competition* Monitored BRET response with varying concentrations of the unlabeled analyte with fixed concentration of the tracer, PTH(1-34)K35<sup>TMR</sup>. *nLuc-ΔECD-PTHR1* Binding of LA-PTH(1-34)K35<sup>TMR</sup> measured with fitting curves into one-phase association model. Binding of M-PTH(1-14)K15<sup>TMR</sup> measured by fitting the curves into one-phase association then dissociation model. All datasets are representative of at least 3 replicates (mean ± SD).

We initially placed the TMR label at the C-terminus of PTH(1-34) or PTHrP(1-34) by appending a Lys residue at position 35 and linking the fluorophore to the Lys side chain. Varying the position of the (TMR)-Lys residue, to position 13 in PTHrP(1-34) or to position 33 in PTH(1-34), did not cause a significant change in the kinetic parameters (Table 1). These observations suggest that our strategy characterizes contacts between the peptide agonist and the PTHR1 with minimal influence from the fluorophore attached to the peptide. The data indicate that PTH(1-34) has modestly higher affinity (∼4-fold) for the PTHR1 relative to PTHrP(1-34).

The relative affinities of human PTH(1-34) and PTHrP(1-34) for the human PTHR1 emerging from our studies are consistent with earlier measurements involving radiotracer displacement. At least three studies have made this comparison based on binding to receptors on whole cells. Gardella et al.^21^ compared human PTH(1-34) and PTHrP(1-34) binding to the rat PTHR1 expressed in two cell types; for both ROS 17/2.8 cells and AR-C40 cells, IC_50_ for PTH(1-34) was approximately two-fold smaller than IC_50_ for PTHrP(1-34). Comparison of PTH(1-34) and PTHrP(1-36) for binding to COS-7 cells expressing the human PTHR1 suggested approximately seven-fold smaller IC_50_ for PTH(1-34).^22^ A comparison of human PTH(1-34) and PTHrP(1-36) for binding to the rat PTHR1 on UMR 106 cells yielded indistinguishable IC_50_ values.^30^

Two studies have employed radiotracer displacement to compare the binding of PTH(1-34) and PTHrP(1-36) to distinct forms of the human PTHR1 in isolated cell membranes. In one version of this assay, the intracellular surface of the PTHR1 is bound to the G_S_ heterotrimer. This form of the receptor is designated the RG state. In the other version of this assay, the PTHR1 is functionally uncoupled from the G protein. This form of the receptor is designated the R^0^ state. Using membranes prepared from COS-7 cells that transiently overexpressed the human PTHR1, Dean et al. reported that human PTH(1-34) and PTHrP(1-36) have similar IC_50_ values for the RG state, but that for the R^0^ state PTH(1-34) displays a ∼4-fold lower IC_50_ relative to PTHrP(1-36).^23^ A similar comparison was subsequently made by Hattersley et al. with membranes from GP2.3 cells, which stably express the human PTHR1.^24^ Again, human PTH(1-34) and PTHrP(1-36) displayed similar IC_50_ values in the RG state assay, but PTH(1-34) bound with higher affinity to the R^0^ state, as indicated by a ∼9-fold lower IC_50_ relative to PTHrP(1-36). Comparison of our results to these precedents suggests that our cell-based assay may reflect the R^0^ state of the PTHR1, as has been predicted for cells that overexpress only the receptor but not G proteins.^25^

Long-acting PTH (LA-PTH) is a designed agonist containing PTH-derived residues in the N-terminal region and PTHrP-derived residues in the C-terminal region.^26^ LA-PTH induces PTHR1 activation of very long duration in cellular assays and in animals.^14^ Binding measurements involving radiotracer displacement indicated that the affinity of LA-PTH for the RG state of the PTHR1 is similar to the RG affinities of PTH(1-34) and PTHrP(1-36), while the affinity of LA-PTH for the R^0^ state is higher relative to the affinities of PTH(1-34) and PTHrP(1-36).^24^ We conducted binding measurements with LA-PTH bearing a TMR unit on a lysine residue at the C-terminus. *k*_*on*_ for this peptide was similar to that of PTH(1-34) or PTHrP(1-34), but *k*_*off*_ for LA-PTH was an order of magnitude lower than *k*_*off*_ for the other two peptides (Table 1). Thus, K_D_ for LA-PTH was smaller than for PTH(1-34) or PTHrP(1-34), which is consistent with the reported trend in IC_50_ values for the R^0^ state.^24^

Ferrandon et al. conducted single-cell FRET studies of agonist binding to a GFP-PTHR1 fusion protein using peptides bearing a TMR fluorophore at position 13.^31^ These measurements, which have a much shorter time scale relative to our BRET measurements, supported a two-state binding mechanism. In this mechanism, the C-terminal portion of the agonist associates with the receptor ECD in the initial step. In a second, slower step, the N-terminal portion of the agonist engages the receptor TMD. For the first step, PTH(1-34) and PTHrP(1-36) displayed *k*_*on*_ values similar to those we measured, but *k*_*off*_ was ∼50-to 100-fold faster for this step than for the interaction characterized in our assay. The two agonists differed dramatically in the second step, according to the single-cell FRET study, with PTHrP(1-36) displaying a *k*_*off*_ value comparable to the value we measured, but little dissociation observed for PTH(1-34).^31^ This latter behavior was attributed to rapid internalization of the PTHR1-PTH(1-34) complex, which continues to stimulate cAMP production from endosomes. In addition to the shorter time scale, the measurements of Ferrandon et al. differ from our BRET-based measurements in that our studies employ cells pretreated with NaN_3_ to prevent receptor internalization, while internalization mechanisms were active under the conditions used by Ferrandon et al.^31^

Overall, the comparisons with previous studies summarized above suggest that the results provided by the new BRET-based assay are consonant with results for agonist-PTHR1 association obtained by other methods. The BRET-based assay can be performed with a conventional luminescence plate reader, making this assay more easily implemented than the assays previously employed to characterize ligand-PTHR1 binding.

### Competition assay format

For comparative studies of multiple peptides, it is convenient to conduct binding assays that involve displacement of a single labeled tracer peptide (Figure 2A). To evaluate the nLuc-PTHR1 construct in competition mode, we used 50 nM PTH(1-34)-K35^TMR^ as the tracer and unlabeled competitors at variable concentration, and monitored tracer binding over time. As the concentration of the competitor peptide was increased, the degree of tracer binding diminished. Figure 2B shows data for tracer binding in the presence of varying concentrations of PTH(1-34), and Figure 2C shows analogous data with PTHrP(1-34). The *k*_*on*_, *k*_*off*_ and K_D_ values derived from these data were similar to those measured with TMR-bearing peptides in the direct-binding format (Table 1).

**Figure 2.**
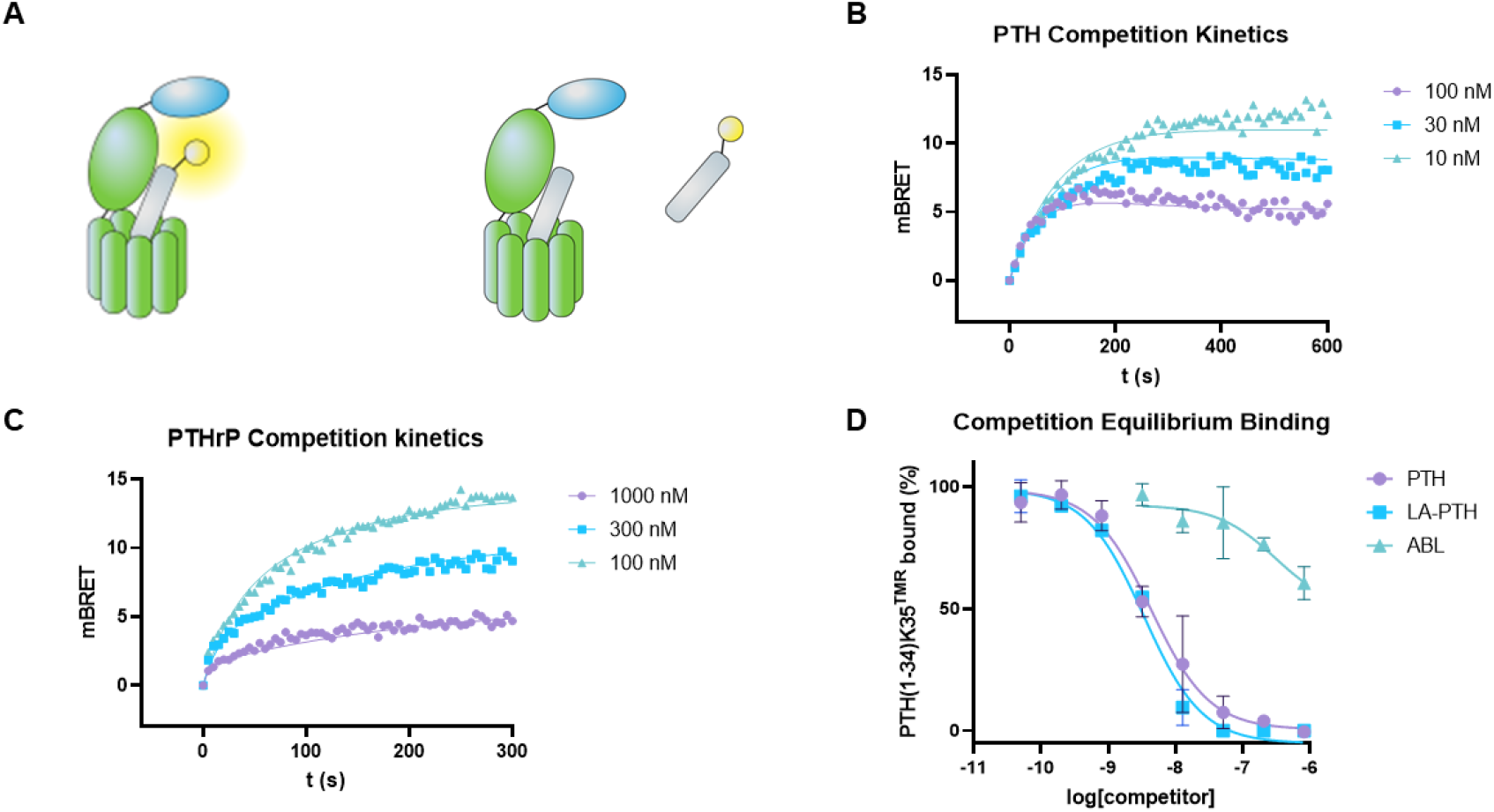
(A) Depiction of competition mode of NanoBRET binding kinetics. nLuc-PTHR1 is introduced to a fixed concentration of the tracer and varying concentrations of competitors. Decrease in the BRET response can be used to calculate the kinetic properties and the affinities of the competitors (B), (C) Competition kinetic binding curves generated using 50 nM PTH(1-34)K35^TMR^ and varying concentrations of PTH(1-34) or PTHrP(1-34). (D) Competition equilibrium binding of PTH, LA-PTH, and ABL against constant 30 nM concentration of PTH(1-34)K35^TMR^.

We used equilibrium competition binding experiments with 30 nM PTH(1-34)-K35^TMR^ as the tracer to evaluate the designed agonist LA-PTH. These measurements suggested IC_50_ of 4.1 ± 1 nM for LA-PTH, compared to 16 ± 10 nM for PTH. Thus, the difference in IC_50_ values between PTH and LA-PTH measured in the competition binding assay (∼4-fold) is comparable to the difference in K_D_ between these two peptides measured in the direct-binding assay (∼5-fold; Table 1). Another PTHR1 agonist, ABL, which is as potent as PTH(1-34) in terms of stimulating cAMP production^24^, failed to fully inhibit PTH(1-34)K35^TMR^ binding even at 1 µM ABL (Figure 2D). This observation is consistent with a previous report that ABL has diminished affinity for the R^0^ state^24^ and our observation that ABL(1-34)K35^TMR^ failed to produce a significant BRET response with nLuc-PTHR1, in contrast to PTH(1-34)K35^TMR^.

Garton et al. reported that a peptide comprised entirely of D-amino acid residues displayed potency comparable to that of PTH(1-34) (L-amino acid residues) in activating the PTHR1.^32^ In our equilibrium binding competition BRET assay, displacement of PTH(1-34)K35^TMR^ by this D-peptide from the receptor was detectable only at the highest peptide concentration employed, 10 µM (Figure S6.). The ability of this D-peptide to activate the PTHR1 was extremely weak in our cAMP assay, with estimated EC_50_ at least 10,000-fold higher than EC_50_ for PTH(1-34).

### Modified assay for binding to a version of the PTHR1 lacking the extracellular domain

Gardella et al. demonstrated that an N-terminally truncated version of the PTHR1 lacking most of the extracellular domain is competent for signal transduction.^27^ PTH(1-34) is a weak agonist of the truncated PTHR1, but other designed peptides, including LA-PTH and M-PTH(1-14), potently activate the truncated receptor. M-PTH(1-14) differs only modestly from the first 14 residues of LA-PTH (Figure 3A). M-PTH(1-14) contains aminoisobutyric acid at positions 1 and 3, which are Ala in LA-PTH. M-PTH(1-14) contains homoarginine at position 11, while LA-PTH contains arginine at this position.

**Figure 3.**
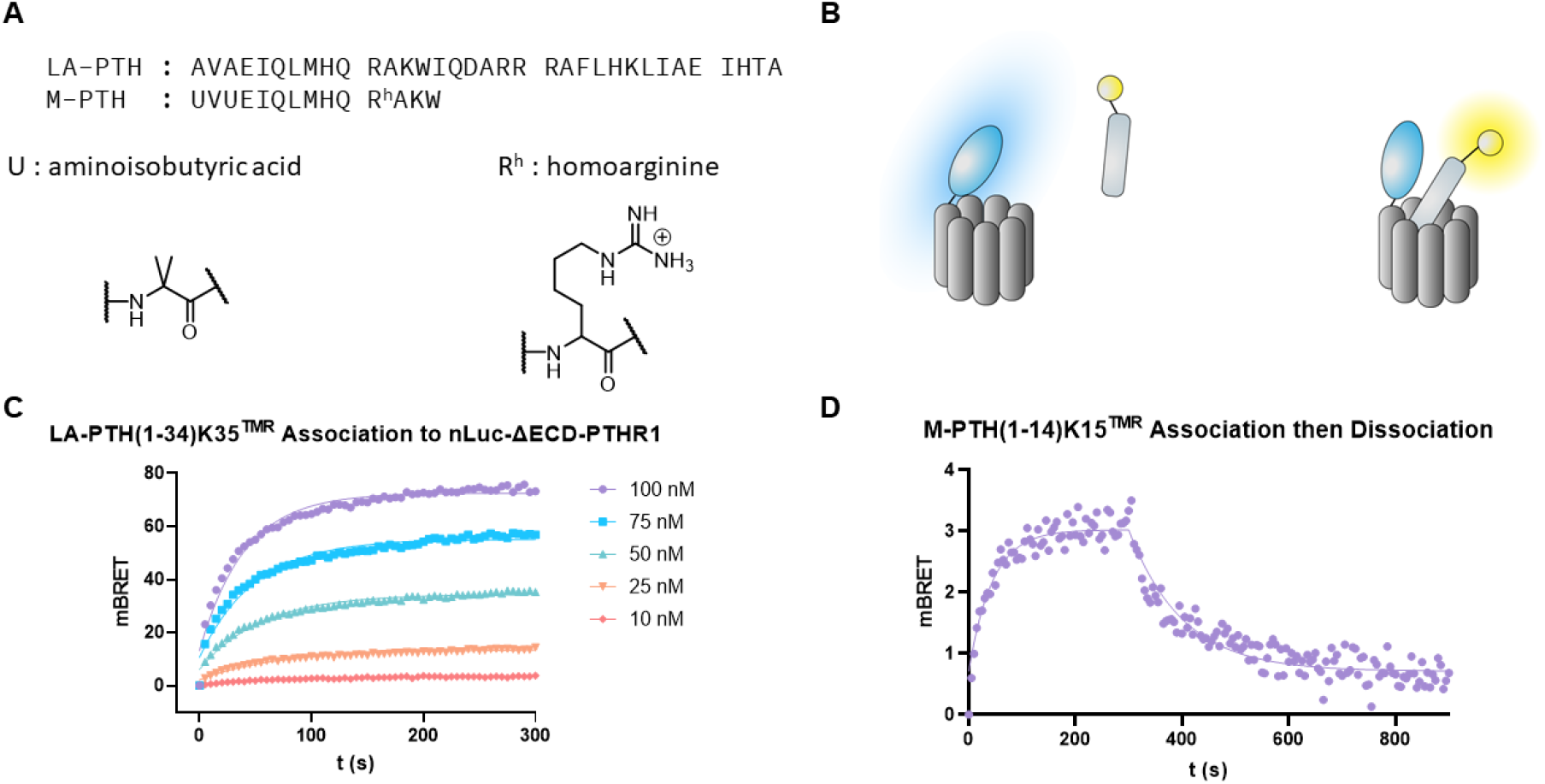
(A) Sequences of LA-PTH(1-34) and M-PTH(1-14). (B) Cartoon depiction of BRET between fluorescently labeled ligand and nLuc-ΔECD-PTHR1 after binding. (C) Association curves of LA-PTH(1-34)K35^TMR^ binding to nLuc-ΔECD-PTHR1. (D) Association then dissociation curves of M-PTH(1-14)K15^TMR^ on nLuc-ΔECD-PTHR1.

We constructed a fusion protein containing the truncated PTHR1 preceded by a nanoluciferase unit (nLuc-ΔECD-PTHR1) (Figure 3B). HEK293H cells transiently transfected with this receptor were used to conduct BRET-based binding studies as described above. Binding of TMR-labeled derivatives of PTH(1-34) and PTHrP(1-34) could not be reliably detected. In each case, a BRET signal was observed at peptide concentrations of 200 nM or higher, but control studies showed that at these TMR-peptide concentrations, BRET or radiative energy transfer can arise without specific binding to the receptor. In contrast, binding could be measured for derivatives of LA-PTH and M-PTH(1-14) bearing a TMR unit (Table 1). For LA-PTH, data for peptide association could be fitted to a pseudo-first order binding model as described above to generate *k*_*on*_ and *k*_*off*_. For binding of M-PTH(1-14)K15^TMR^, however, we employed an association-then-dissociation approach. *k*_*off*_ was calculated from the dissociation curve based on a one-phase exponential decay model, and this rate constant was then used to fit the association curve and calculate the *k*_*on*_ value. *k*_*on*_ measured for LA-PTH with the nLuc-ΔECD-PTHR1 was indistinguishable from *k*_*on*_ measured for this peptide with the full-length nLuc-PTHR1, but *k*_*off*_ measured for the truncated receptor was considerably larger relative to *k*_*off*_ for the full-length receptor. The similarity of the *k*_*on*_ values suggests that this rate constant is primarily determined by interaction between LA-PTH and the receptor TMD, even when the ECD is present. This conclusion is consistent with the recent cryo-EM structure for the complex between LA-PTH and the PTHR1.^13^ Three bound states were resolved among the particles, all with approximately the same interactions between the N-terminal portion of LA-PTH and the receptor TMD. However, significant variations were observed in the relationship between the C-terminal portion of LA-PTH and the ECD, with one of the three states showing no interaction in this region of the complex. The difference in *k*_*off*_ values at the full-length receptor for LA-PTH vs. M-PTH(1-14) suggests that interactions between the C-terminal portion of LA-PTH and the ECD contribute significantly to the overall affinity of the receptor for LA-PTH.

### Sequence variants of known PTHR1 agonists

We selected several peptides derived from PTH(1-34) and/or PTHrP(1-36) for evaluation with the new assay based on previous reports documenting effects of sequence modifications on agonist activity and/or PTHR1 binding. The goals of these studies were: (1) to enhance our understanding of the relationship between data generated by the new assay and previous measurements, and (2) to provide new insights on previously described agonists.

#### N-terminal truncations of PTH(1-34)

Early studies with PTH(1-38) showed that bone anabolic effects in rats were diminished upon removal of the N-terminal residue (to generate PTH(2-38)), and these effects were lost altogether with a further truncation (to generate PTH(3-38)).^33^ We observed that PTH(2-34) was moderately less potent than PTH(1-34), as indicated by a ∼10-fold increase in EC_50_ for PTH(2-34) relative to PTH(1-34), while PTH(3-34) showed only minimal activity (Figure 4A). Derivatives of these N-terminally truncated peptides containing Lys at the C-terminus with TMR on the side chain were very similar to the peptides lacking TMR in terms of their ability to stimulate intracellular cAMP production (Figure S7). In studies with the nLuc-PTHR1, *k*_*on*_ values for PTH(2-34)K35^TMR^ and PTH(3-34)K35^TMR^ (Figure 4B,C) were indistinguishable from that of the full-length analogue, PTH(1-34)K35^TMR^ (Table 2). The *k*_*off*_ values were indistinguishable for the full-length peptide and PTH(2-34)K35^TMR^, while *k*_*off*_ was modestly larger for PTH(3-34)K35^TMR^ (Table 2). These results show that although the N-terminal residues of PTH are critical in terms of receptor activation, these two residues contribute little or nothing to peptide affinity for the PTHR1. The recent cryo-EM structure of the complex between LA-PTH and the PTHR1 suggests that contacts between the N-terminus of the peptide and residues in receptor TM helix 6 promote adoption of the activated conformation of the PTHR1.^13^ It has been proposed that these contacts are repulsive,^34^ a view that seems to be consistent with our binding data.

**Table 2.**
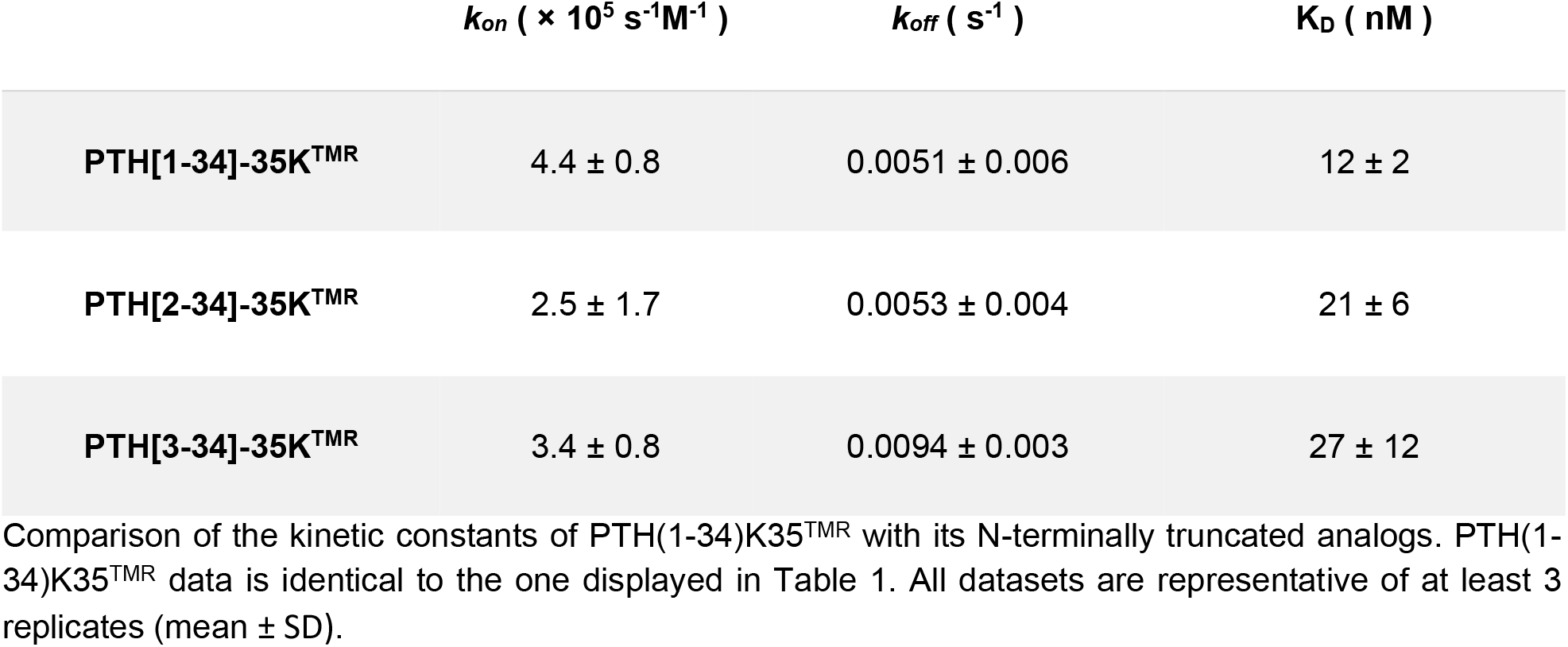
The kinetic constants and dissociation constants of two N-terminally truncated analogs of PTH.

**Figure 4.**
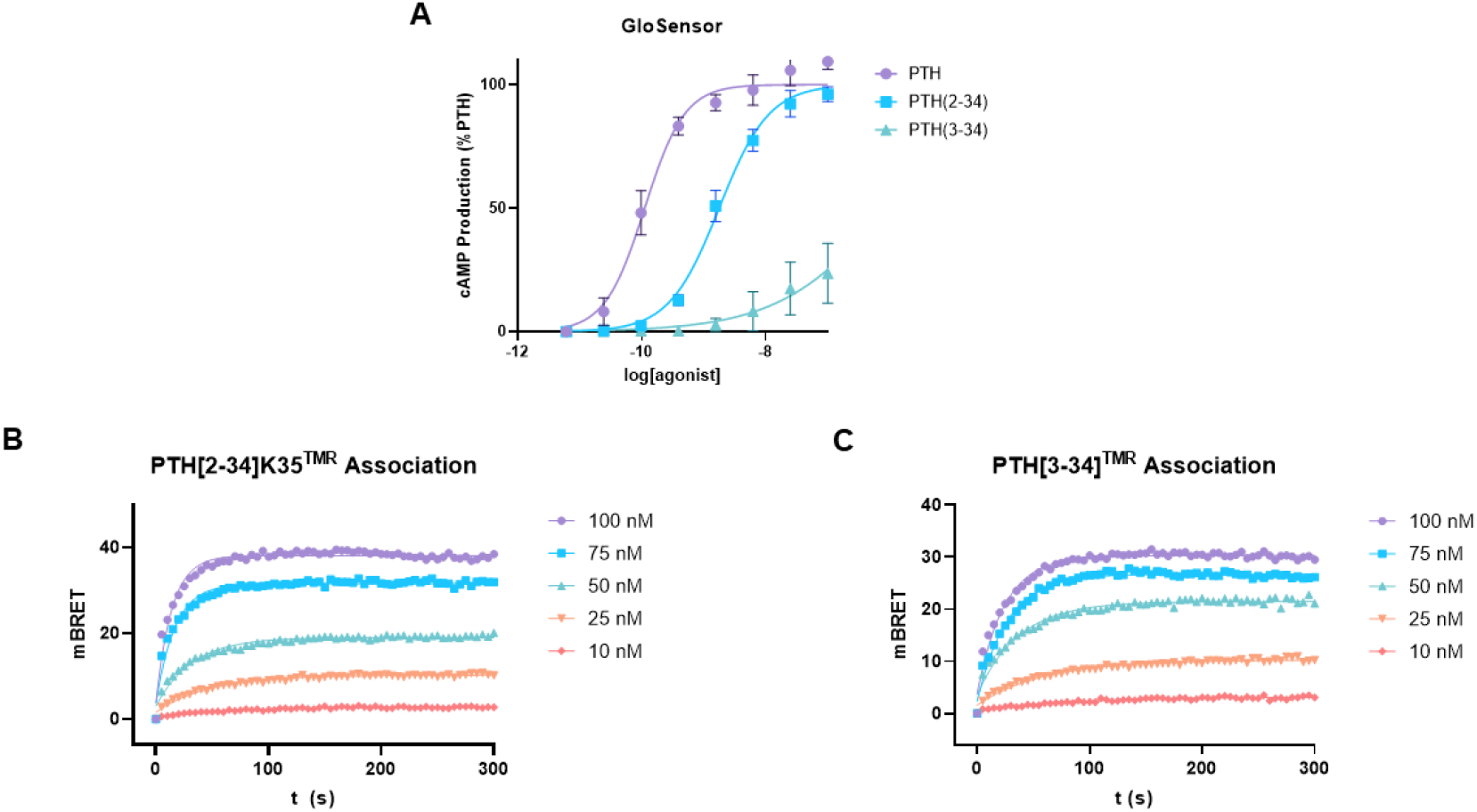
(A) cAMP activities of PTH(1-34) and its N-terminally truncated analogs, PTH(2-34) and PTH(3-34). (B), (C) Association binding curves of PTH(2-34)K35^TMR^ and PTH(3-34)K35^TMR^ on nLuc-PTHR1.

#### Variations at position 5 of PTH and PTHrP

There are many differences between the sequences of PTH(1-34) and PTHrP(1-34), but the difference at position 5 (Ile in PTH vs. His in PTHrP) is known to have a particularly large effect in terms of affinity for the receptor and signaling profile.^21,22^ Comparisons involving PTH(1-34) and the I5H variant, and PTHrP(1-36) and the H5I variant, revealed that neither change caused a substantial alteration in affinity for the RG state of the PTHR1.^23^ In contrast, these changes had large effects on affinity for the R^0^ state, with Ile at position 5 being favored in both the PTH and PTHrP contexts. Thus, native PTH(1-34) displayed ∼20-fold lower IC_50_ relative to the I5H variant in the competition binding assay for the R^0^ state, and the H5I variant of PTHrP(1-36) displayed a ∼10-fold lower IC_50_ relative to the version with the native His. These binding assays were conducted in DPBS buffer at pH 7.4.^23^

We used the BRET assay (direct binding mode) to compare PTH(1-34)K35^TMR^ with the I5H variant, and to compare PTHrP(1-34)K35^TMR^ with the H5I variant (Table 3). Trends from measurements conducted at pH 7.5 were qualitatively consistent with those reported for R^0^ state binding based on radioligand displacement.^23^ For PTH(1-34), the native Ile5 led to ∼6-fold lower K_D_ relative to non-native His5, while for PTHrP(1-36), the non-native Ile5 led to ∼5-fold lower K_D_ relative to native His5. The major kinetic effect of Ile vs. His at position 5 was manifested in *k*_*off*_, which was an order of magnitude larger with His in both cases. *k*_*on*_ was larger with His as well, but only slightly.

**Table 3.** The effect of pH on the kinetic constants and the dissociation constants of PTH(1-34) and PTHrP(1-36), as well as their analogs bearing substitutions on the 5^th^ position.

| | pH | $k_{on} (\times 10^5 \text{ s}^{-1}\text{M}^{-1})$ | $k_{off} (\text{s}^{-1})$ | $K_D (\text{nM})$ |
| --- | --- | --- | --- | --- |
| PTH | 7.5 | $4.8 \pm 0.5$ | $0.0049 \pm 0.002$ | $10 \pm 3$ |
| | 6.5 | $4.2 \pm 0.8$ | $0.0049 \pm 0.001$ | $12 \pm 3$ |
| | 5.5 | $9.8 \pm 4$ | $0.039 \pm 0.006$ | $39 \pm 6$ |
| PTH(I5H) | 7.5 | $7.6 \pm 1$ | $0.049 \pm 0.01$ | $64 \pm 10$ |
| | 6.5 | $6.8 \pm 3$ | $0.071 \pm 0.04$ | $104 \pm 50$ |
| | 5.5 | $4.4 \pm 2$ | $0.088 \pm 0.04$ | $200 \pm 110$ |
| PTHrP | 7.5 | $3.1 \pm 0.5$ | $0.021 \pm 0.006$ | $68 \pm 20$ |
| | 6.5 | $3.4 \pm 0.2$ | $0.049 \pm 0.02$ | $140 \pm 50$ |
| | 5.5 | $2.2 \pm 0.5$ | $0.071 \pm 0.04$ | $330 \pm 160$ |
| PTHrP(H5I) | 7.5 | $2.9 \pm 0.2$ | $0.0035 \pm 0.0005$ | $12 \pm 2$ |
| | 6.5 | $2.7 \pm 0.3$ | $0.0034 \pm 0.0006$ | $12 \pm 2$ |
| | 5.5 | $1.5 \pm 0.3$ | $0.071 \pm 0.02$ | $48 \pm 10$ |

Kinetic constants of PTH(1-34)K35<sup>TMR</sup>, PTHrP(1-36)K35<sup>TMR</sup>, and analogues modified at position 5 at pH 7.5, 6.5, and 5.5. The buffers were prepared with 137 mM NaCl, 1 mM CaCl<sub>2</sub>, 0.5 mM MgCl<sub>2</sub>, 0.02% NaN<sub>3</sub> (w/v), and 0.1% BSA (w/v), using HEPES (7.5), MES (6.5), or acetate (5.5). All datasets are representative of at least 3 replicates (mean ± SD).

The pH 7.5 parameters in Table 3 for PTH(1-34)K35_^TMR^_ and PTHrP(1-36)K35^TMR^ differ slightly from those in Table 1. These small differences may arise because of differences in media for the two assays. The assays summarized in Table 1 were conducted in Dulbecco’s phosphate-buffered saline. We measured the pH of this medium to be in the range 7.0-7.3, which is slightly lower than the HEPES buffer used for the pH 7.5 measurements summarized in Table 1.

The complex formed between the PTHR1 and PTH(1-34) remains intact and continues to stimulate cAMP production after internalization.^31^ In contrast, internalization of the complex formed between PTHrP(1-36) and PTHR1 causes dissociation of the peptide and halts cAMP production.^31^ The divergent behaviors of these two agonists may arise from changes in pH encountered by the peptide-receptor complex as it is trafficked from the cell surface to endosomes.^35^ The shift from neutral pH at the cell surface to the mildly acidic environment of a late endosome (pH ∼ 5.5) should enhance protonation of His side chains. Protonation of His5 of PTHrP(1-36), which is buried in the receptor TMD core, could cause a substantial decline in affinity for the PTHR1. In contrast, protonation at position 5 of PTH(1-34) is not possible because the Ile side chain does not contain a basic site. Thus, the difference between PTH and PTHrP at position 5 could play a critical role in the divergent spatiotemporal signaling profiles of these agonists.

We evaluated binding to the nLuc-PTHR1 as a function of pH (7.5 vs. 6.5 vs. 5.5) for PTH(1-34)K35^TMR^, PTHrP(1-36)K35^TMR^ and the TMR-bearing analogues modified at position 5 to test the hypothesis in the preceding paragraph, i.e., to determine whether a drop in pH causes a larger decline in affinity for the peptides with His at position 5 relative to those with Ile at this position (Table 3). For all four peptides, affinity was modestly lower at pH 5.5 relative to pH 7.5. However, the effect of diminished pH was similar for peptides containing Ile5 and those containing His5. Thus, at pH 5.5 relative to pH 7.5, K_D_ was ∼4-fold larger for PTH(1-34), and K_D_ was ∼5-fold larger for PTHrP(1-36). A comparable change was observed for I5H variant of PTH(1-34). Among the two kinetic parameters, the larger changes were observed in *k*_*off*_, and pH-dependent variation seemed larger for the peptides containing Ile relative to those containing His at position 5. Overall, these data indicate that protonation of the His residue near the N-terminus of PTHrP(1-36) does not constitute a switch that dramatically lowers the affinity of this agonist for the PTHR1 within endosomes compared to receptor at the cell surface.

Both PTHrP(1-34) and PTHrP(1-36) have been subjects of previous studies. To ask whether this small difference in length affects agonism trends, we compared the H5I variants of PTHrP(1-36) and PTHrP(1-34) by measuring affinities for the nLuc-PTHR1 at different pH. Values obtained for PTHrP(1-34)-His5-Ile-K35^TMR^, K_D_ = 5.0 nM at pH 7.5, K_D_ = 5.7 nM at pH 6.5, and K_D_ = 56 nM at pH 5.5, are similar to those obtained for PTHrP(1-36)-His5-Ile-K35^TMR^ (Table 3).

#### PTH/PTHrP hybrids

Early studies evaluated synthetic hybrids of PTH(1-34) and PTHrP(1-34) in terms of binding to and activation of the PTHR1.^21,22^ Binding was assessed via radioligand displacement assays involving whole cells. These studies were motivated by the hypothesis that the N-terminal and C-terminal portions of these peptides likely engaged distinct portions of the receptor, the TMD and ECD, respectively. This hypothesis was subsequently validated by structural analysis of receptor-peptide complexes.^13,14,36–38^

The hybrid PTH(1-14)/PTHrP(15-34) was reported to be indistinguishable from PTH(1-34) in competition binding measurements with the rat PTHR1 expressed in ROS17/2.8 or AR-C40 cells,^21^ but a subsequent comparison involving the human PTHR1 expressed in COS-7 cells indicated weaker binding (∼12-fold higher IC_50_) for the hybrid peptide relative to PTH(1-34).^22^ The complementary hybrid, PTHrP(1-14)/PTH(15-34), in contrast, bound much more weakly than PTH(1-34) in all three comparisons. IC_50_ for the second hybrid was ∼1000-fold larger in the study with ROS17/2.8 or AR-C40 cells and ∼200-fold larger in the study with COS-7 cells. The dramatic difference between the two hybrids in terms of PTHR1 affinity motivated the development of LA-PTH, the C-terminal portion of which is very similar to PTHrP(15-36), and the N-terminal portion of which is derived from PTH(1-14).^26^ In the COS-7 cell study, agonist activities were evaluated.^22^ Despite the large differences in binding, as reflected in IC_50_ values, PTH(1-34), PTHrP(1-36) and the two hybrid peptides were very similar in terms of their ability to stimulate cAMP production, as indicated by EC_50_ and E_max_ values.

These precedents motivated us to evaluate two sets of PTH/PTHrP hybrid peptides, with varying sequence proportions drawn from the two prototype peptides (Table 4). One set contained PTH residues at the N-terminus and PTHrP residues at the C-terminus; the other set had the opposite arrangement. Each peptide had Lys at position 35, with TMR on the side chain. All of these peptides were potent as agonists: EC_50_ for cAMP production varied between 0.1 and 1.0 nM, and each hybrid matched E_max_ achieved with PTH(1-34) (Table 4). This behavior is consistent with the activities reported for hybrid peptides with COS-7 cells.^22^

**Table 4.**
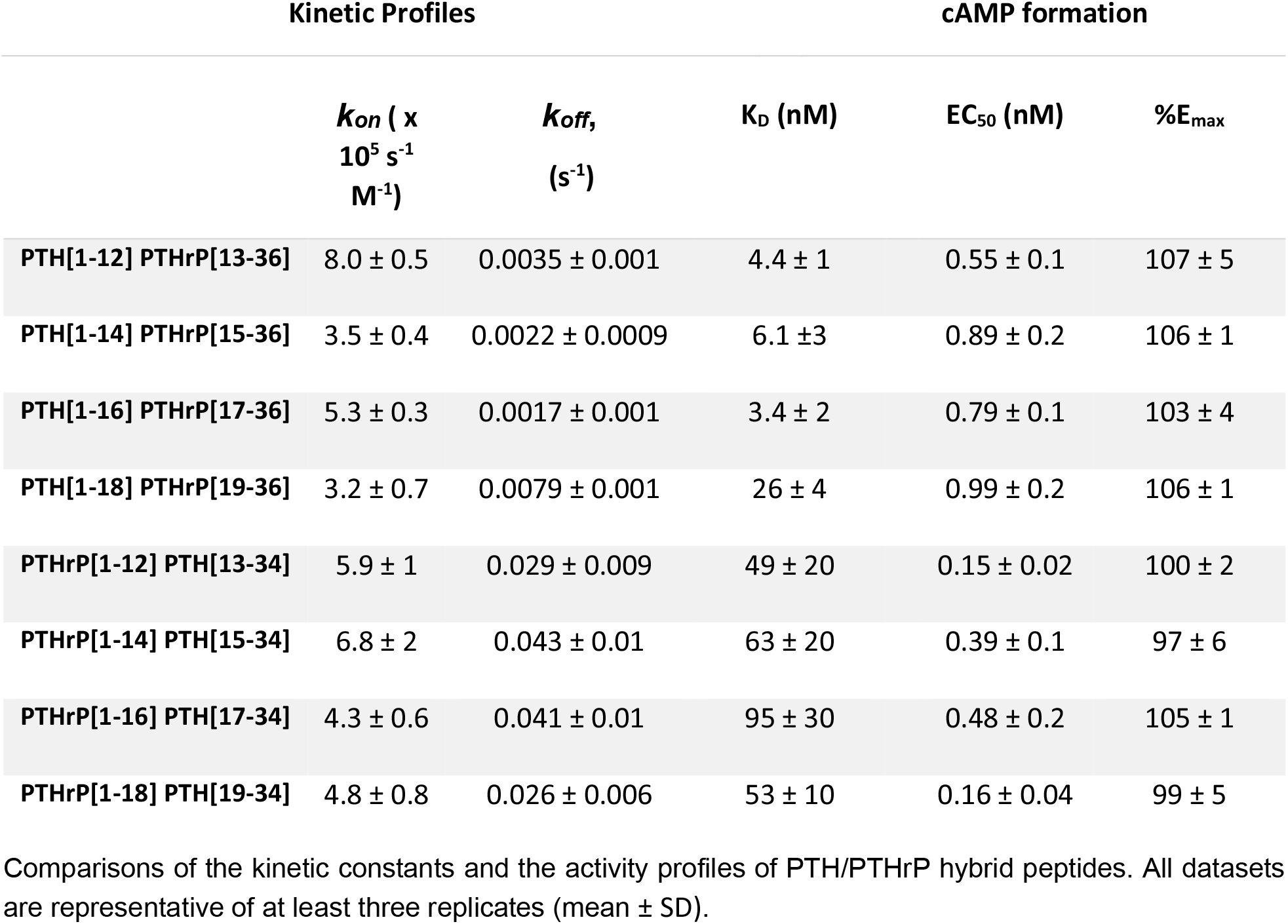
The kinetic constants, dissociation constants, and the activity profiles of various PTH/PTHrP hybrids.

Trends in PTHR1 affinity we measured for the hybrid peptides were qualitatively consistent with those observed in radioligand displacement assays,^22^ although the range of variation in our binding assay was much smaller than in the earlier studies.^9^ The hybrid peptides with N-terminal PTHrP sequence and C-terminal PTH sequence had K_D_ values about one order of magnitude higher than those with N-terminal PTH sequence and C-terminal PTHrP sequence. The point at which the sequence switch occurred had little effect on K_D_ (∼2-fold variation) among the series spanned by PTHrP(1-12)/PTH(13-34) and PTHrP(1-18)/PTH(19-34). A similar trend was observed among PTH(1-12)/PTHrP(13-36), PTH(1-14)/PTHrP(15-36) and PTH(1-16)/PTHrP(17-36), but K_D_ was modestly higher for PTH(1-18)/PTHrP(19-36). *k*_*on*_ values were very similar among all of the hybrid peptides. In contrast, the series with N-terminal PTHrP sequence/C-terminal PTH showed significantly larger *k*_*off*_ values relative to the series with N-terminal PTH sequence/C-terminal PTHrP sequence.

#### Conclusions

We have developed a convenient BRET-based assay for characterizing the binding of agonist peptides to the PTHR1, a target of multiple drugs in current clinical use. This assay format offers practical advantages relative to radioligand displacement. Multiple comparisons with earlier radioligand displacement studies suggest that the BRET assay provides generally consistent results, although some variations were evident.

The BRET assay can be conducted in a “direct binding” mode with agonists that bear a TMR appendage, which is straightforward to introduce in conjunction with solid-phase peptide synthesis. Addition of a TMR group represents a modest change in the context of a 34-to 36-residue peptide, which led us to hypothesize that this modification would not cause a profound alteration in peptide-receptor interactions if the TMR unit were strategically placed. Our data support this hypothesis. In addition, our data indicate that the receptor does not experience a major functional alteration as a result of the fused nanoluciferase domain. We note that the BRET-based strategy has been applied recently to family A GPCRs, the agonists of which are similar in size to the fluorophore that must be attached. In such cases, it is easy to imagine that fluorophore attachment could exert substantial effects on agonist performance.

Use of peptides bearing a TMR group in the BRET assay allows direct measurement of the rate constants *k*_*on*_ and *k*_*off*_ for agonist-PTHR1 interaction, and K_D_ can be determined from those values. Alternatively, the BRET assay can be run in competition format, which accommodates unlabeled peptides and streamlines comparisons.

We have used the new assay to ask whether the Ile vs. His difference at position 5 of PTH and PTHrP underlies the difference in spatiotemporal signaling profiles between these two agonists. PTH continues to stimulate cAMP production after the agonist-receptor complex has been trafficked to endosomes, while PTHrP signaling is terminated upon endocytosis.^31^ There is great interest in PTHR1 signaling timeframe because the temporal pattern of agonist administration strongly influences the impact on bone growth.^39,40^ Our data suggest that endosomal acidification does not lead to a large change in the relative affinities of PTH(1-34) and PTHrP(1-36), as would have been predicted if His protonation destabilized the bound form of PTHrP relative to the bound form of PTH.

The new assay provides insights that complement early studies of N-terminal truncation of PTH-derived peptides.^33^ Removal of the first residue substantially impairs receptor activation, as manifested in stimulation of cAMP production, and removal of the first two residues further hinders agonist activity. Measurements from the BRET assay, however, show that PTH(2-34) and PTH(3-34) are comparable to PTH(1-34) in terms of affinity for the PTHR1. On the other hand, use of the BRET assay to examine a D-peptide^32^ that was reported to be comparable to PTH(1-34) in potency as an agonist for the PTHR1 indicates that the D-peptide binds only very weakly to the receptor. These observations prompted us to evaluate the D-peptide for the ability to stimulate cAMP production via the PTHR1; only very weak activity was detected, which is inconsistent with the earlier report.

The new BRET assay based on the nLuc-PTHR1 fusion should be useful to researchers who study signal transduction mediated by this medically important receptor. The construct will be available via Addgene, which should facilitate exploration by others.

## Methods

### Molecular Biology

Nanoluciferase was fused to the N-terminus of the PTHR1 using Gibson assembly. Briefly, plasmids encoding nLuc-GLP-1R^41^ and Human PTHR1 in a pcDNA3.1+ vector (cDNA Resource Center, #PTHR100000) were linearized and amplified with appropriate overhangs to facilitate Gibson assembly using the Phusion HF polymerase PCR kit (Thermo) according to the manufacturer’s instructions. The PCR products were treated with DpnI (0.8 μl; Agilent) for 1 h at 37 °C, and DNA was isolated using concentrator spin columns (Zymo). Then, 0.03 pmol and 0.06 pmol of linearized DNA encoding receptor + vector and NLuc, respectively, were assembled using 10 μl of 2× master mix at 50 °C for 1 h according to manufacturer’s instructions (NEB). 5α Competent *Escherichia coli* (30 μl; NEB) was then transformed with 2 μl of the assembly reaction mixture by heat shocking for 30 s at 42 °C. These cells were then allowed to grow in SOC medium (0.5 ml) for 1 h at 37 °C with shaking. The cell suspension was spread onto LB agar plates with ampicillin (100 μg ml^-1^), the plates were incubated overnight at 37 °C, and colonies were selected for plasmid Maxiprep (Qiagen) according to the manufacturer’s protocol. Site-directed mutagenesis using the QuikChange II XL Site-Directed Mutagenesis Kit (Agilent) according to the manufacturer’s instructions was used to install the desired stop codon. Sanger sequencing performed at the University of Wisconsin-Madison Biotechnology Center DNA Sequencing Facility was used to confirm the desired sequence.

### Peptide synthesis

Peptides were prepared by microwave-assisted solid-phase peptide synthesis using either a CEM Mars microwave reactor (manual) or an automated CEM Liberty Blue reactor. Low-loading Rink Amide resin (CEM) was used as the solid support. For peptide coupling steps, 5 equiv. of Fmoc-amino acid, 10 equiv. of diisopropylcarbodiimide, and 5 equiv. of Oxyma with respect to the resin loading were used. NMP or DMF solutions of these compounds were added to the reaction vessel. The resulting slurry was heated to 90°C and stirred for 2 min, except for histidine, which was heated to 50°C for 10-30 min. After the heating, the resin was washed with DMF thrice, and the Fmoc was removed by adding 20% piperidine solution in DMF with 0.1% Oxyma, and heating to 80°C for 2 min. For orthogonal removal of the alloc protecting group on a lysine side chain, the resin was washed with methylene chloride thrice, and 10 equiv. of phenylsilane and 1 equiv. of tetrakis(triphenylphosphine)palladium(0) in ethylene dichloride were added to the reaction vessel. The resulting slurry was heated to 35°C for 10 min. This process was repeated twice. The resin was washed with methylene chloride thrice and dimethylformamide thrice. 4 equiv. 6-caboxytetramethylrhodamine, 4 equiv. of diisopropylcarbodiimide, and 4 equiv. of Oxyma in biotech-grade DMF were added, and the resulting mixture was heated to 75°C for 10 min. At the end of the solid-phase synthesis, the peptide was cleaved and deprotected in a solution containing 90% TFA, 5% thioanisole, 3% DODT and 2% anisole by incubating at room temperature for 4 h. The resulting solution was filtered from the resin, and product was precipitated by addition of 10-fold excess of −20 °C diethyl ether. Collected precipitate was washed twice with diethyl ether, dried, and dissolved in DMSO for further purification with a preparative reverse-phase HPLC system (Waters) using a C-18 column (12 mL/min flowrate, with gradients of H_2_O + 0.1% TFA as solvent A and MeCN + 0.1% TFA as solvent B). The collected fractions were analyzed based on mass using MALDI-TOF-MS (Bruker UltraFlex) with CHCA as matrix. Isolated peptide was assessed for purity using a UPLC system (Waters) with a C-18 column (0.3 mL/min flowrate, with gradients of H_2_O + 0.1% TFA as solvent A and MeCN + 0.1% TFA as solvent B). The concentration of peptide in stock solutions was determined using a NanoDrop One UV/Vis system (Thermo Scientific).

### Cell Culture

Cells were grown in Dulbecco’s Modified Eagle Medium (with D-glucose but lacking L-glutamine and sodium pyruvate) supplemented with 10% fetal bovine serum (Corning, Cat# 45000-734). Cells were incubated at 37°C with 5% CO_2_. Dulbecco’s Phosphate Buffered Saline (DPBS; Gibco, Cat#14190235) and trypsin-EDTA (Corning, Cat# 45001-082) at 0.05% were used to detach cells for passaging and transfer to assay plates.

### Transfection

HEK293FT cells were seeded onto a 100 mm dish (Fisher, Cat# FB012924) and grown to 80-90% confluency as observed under a microscope. On the day of transfection, 10 µg of plasmid was mixed with 30 µL of FugeneHD (Promega, Cat# PAE2311) in Opti-MEM (Gibco, Cat# 31985062) to bring the final volume to 1 mL. The resulting mixture was gently mixed by flicking the Eppendorf tube,which was set to incubate at room temperature for 20-30 min. The medium from the 100 mm dish was aspirated, and the cells were washed with DPBS without calcium or magnesium. To the cells was added 4.5 mL of McCoy’s 5A supplemented with 10% FBS, and the cells were incubated at 37°C for 5 min. Following the incubation, 4.5 mL of DMEM with 10% FBS was added, and the transfection solution was added gently over the cells. The cells were left to incubate at 37°C for 24-36 h.

### Glosensor Assay

HEK293 cells stably expressing the WT-PTHR1 and the Glosensor construct (GP2.3) were grown in DMEM with 10% FBS. GP2.3 cells were harvested and seeded onto a 96-well assay plate (Costar, Cat# 29444-041), which was incubated at 37°C. On the day of the experiment, the medium was aspirated, and 90 µL of DPBS solution containing 500 µM of D-luciferin was added to each well. The cells were incubated at room temperature for 20 min. To the cells was added 10 µL of the peptide solution at various concentrations, and the luminescence was measured using a Biotek plate reader over 30 min. The luminescence at the 15-minute time point was used for activity analysis.

### Kinetics Assay using BRET

HEK293H cells stably expressing the nLuc-PTHR1 were generated using zeocine resistance as the selective marker. The cells were grown in DMEM supplemented with 10% FBS. Before the day of the experiment, the cells were seeded onto a white opaque 96-well assay plate at 5,000-10,000 cells per well. The cells were incubated at 37°C for 24 h. On the day of the experiment, the medium was aspirated, and 70 µL of DPBS supplemented with 0.02% NaN_3_, 1 mM CaCl_2_ and 0.5 mM MgCl_2_ was added. The cells were incubated at room temperature for at least 30 min before the experiment. The labeled peptide solution was prepared in the same solution that also included 5 µM h-coelenterazine. For the experiment, the medium was aspirated from the well, 100 µL of the peptide solution with h-coelenterazine was added to the well, and the luminescence from 460 nm and 590 nm was measured over 5-10 min at 5-second intervals. The ratio of the luminescence from the 590 nm channel over that from the 460 nm channel with a 1,000x multiplier was used as the mBRET value for generating plots. For nLuc-ΔECD-PTHR1, HEK293FT cells were transiently transfected with the appropriate construct.

### Competition Binding Kinetics

The general procedure for preparation was the same as for the direct binding assay. For the assay buffer, 50 nM PTH(1-34)K35^TMR^ and three concentrations of each unlabeled ligand were prepared; each solution contained 5 µM h-coelenterazine. After ligand addition, the three wells (one for each unlabeled ligand concentration) were monitored at 9-second intervals.

### Data Analysis

For the direct binding kinetics experiments, the mBRET values were plotted against time in seconds, and fitted to equation 1. The extracted *k*_*obs*_ values were then plotted against the concentration of the tracer in equation 2 to calculate the slope and the Y-intercept, corresponding to *k*_*on*_ and *k*_*off*_, respectively. The K_D_ values were calculated as the ratio of *k*_*on*_ and *k*_*off*_ (equation 3). For the association-then-dissociation kinetics experiment, the dissociation curve was fitted to a one-phase dissociation with Graphpad Prism software to measure *k*_*off*_. Then the *k*_*off*_ value was used as a constraint for fitting the association curve to extract the *k*_*on*_ value. For the competition kinetics experiment, the curves generated were fitted into “Competition Kinetics” in Graphpad Prism software with the predetermined *k*_*on*_, *k*_*off*_ and concentration values of the tracer to calculate the *k*_*on*_ and *k*_*off*_ values of the unlabeled peptide.

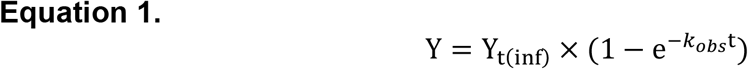

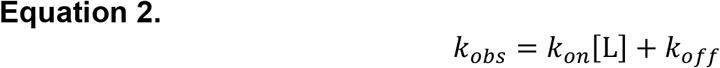

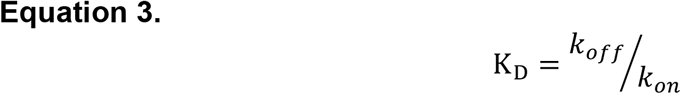

## Acknowledgements

This work was supported in part by NIH grant R01 GM056414. B.P.C. was supported in part by a graduate fellowship from the NSF (DGE-1747503) and by a Biotechnology Training Grant from NIH (T32 GM008349). K.D.N. was supported by a Hilldale Fellowship from UW-Madison.

## Notes

### Competing Interest Statement

The authors have declared no competing interest.

## References

(1) Gardella, T. J.; Vilardaga, J. The Parathyroid Hormone Receptors — Family B G Protein – Coupled Receptors. Pharmacological Reviews 2015, 67 (2), 310–337. https://doi.org/10.1124/pr.114.009464.

(2) Cheloha, R. W.; Gellman, S. H.; Vilardaga, J. P.; Gardella, T. J. PTH Receptor-1 Signalling - Mechanistic Insights and Therapeutic Prospects. Nature Reviews Endocrinology 2015, 11 (12), 712–724. https://doi.org/10.1038/nrendo.2015.139.

(3) Sutkeviciute, I.; Clark, L. J.; White, A. D.; Gardella, T. J.; Vilardaga, J. P. PTH/PTHrP Receptor Signaling, Allostery, and Structures. Trends in Endocrinology and Metabolism 2019, 30 (11), 860–874. https://doi.org/10.1016/j.tem.2019.07.011.

(4) Blick, S. K. A.; Dhillon, S.; Keam, S. J. Teriparatide. Drugs 2008, 68 (18), 2709–2737. https://doi.org/10.2165/0003495-200868180-00012.

(5) Shirley, M. Abaloparatide: First Global Approval. Drugs 2017, 77 (12), 1363–1368. https://doi.org/10.1007/s40265-017-0780-7.

(6) Mannstadt, M.; Bilezikian, J. P.; Thakker, R. v.; Hannan, F. M.; Clarke, B. L.; Reijnmark, L.; Mitchell, D. M.; Vokes, T. J.; Winer, K. K.; Shoback, D. M. Hypoparathyroidism. Nature Reviews Disease Primers 2017, 3 (17055), 1–20. https://doi.org/10.1038/nrdp.2017.55.

(7) Thomas, B. E.; Woznica, I.; Mierke, D. F.; Wittelsberger, A.; Rosenblatt, M. Conformational Changes in the Parathyroid Hormone Receptor Associated with Activation by Agonist. Molecular Endocrinology 2008, 22 (5), 1154–1162. https://doi.org/10.1210/me.2007-0520.

(8) Wootten, D.; Miller, L. J.; Koole, C.; Christopoulos, A.; Sexton, P. M. Allostery and Biased Agonism at Class b g Protein-Coupled Receptors. Chemical Reviews 2017, 117 (1), 111–138. https://doi.org/10.1021/acs.chemrev.6b00049.

(9) Bortolato, A.; Doré, A. S.; Hollenstein, K.; Tehan, B. G.; Mason, J. S.; Marshall, F. H. Structure of Class B GPCRs: New Horizons for Drug Discovery. British Journal of Pharmacology 2014, 171 (13), 3132–3145. https://doi.org/10.1111/bph.12689.

(10) Hollenstein, K.; de Graaf, C.; Bortolato, A.; Wang, M. W.; Marshall, F. H.; Stevens, R. C. Insights into the Structure of Class B GPCRs. Trends in Pharmacological Sciences 2014, 35 (1), 12–22. https://doi.org/10.1016/j.tips.2013.11.001.

(11) Liang, Y. L.; Belousoff, M. J.; Zhao, P.; Koole, C.; Fletcher, M. M.; Truong, T. T.; Julita, V.; Christopoulos, G.; Xu, H. E.; Zhang, Y.; Khoshouei, M.; Christopoulos, A.; Danev, R.; Sexton, P. M.; Wootten, D. Toward a Structural Understanding of Class B GPCR Peptide Binding and Activation. Molecular Cell 2020, 77 (3), 656-668.e5. https://doi.org/10.1016/j.molcel.2020.01.012.

(12) Cong, Z.; Liang, Y. L.; Zhou, Q.; Darbalaei, S.; Zhao, F.; Feng, W.; Zhao, L.; Xu, H. E.; Yang, D.; Wang, M. W. Structural Perspective of Class B1 GPCR Signaling. Trends in Pharmacological Sciences 2022, 43 (4), 321–334. https://doi.org/10.1016/j.tips.2022.01.002.

(13) Zhao, L. H.; Ma, S.; Sutkeviciute, I.; Shen, D. D.; Edward Zhou, X.; de Waal, P. W.; Li, C. Y.; Kang, Y.; Clark, L. J.; Jean-Alphonse, F. G.; White, A. D.; Yang, D.; Dai, A.; Cai, X.; Chen, J.; Li, C.; Jiang, Y.; Watanabe, T.; Gardella, T. J.; Melcher, K.; Wang, M. W.; Vilardaga, J. P.; Eric Xu, H.; Zhang, Y. Structure and Dynamics of the Active Human Parathyroid Hormone Receptor-1. Science (1979) 2019, 364 (6436), 148–153. https://doi.org/10.1126/science.aav7942.

(14) Ehrenmann, J.; Schöppe, J.; Klenk, C.; Rappas, M.; Kummer, L.; Doré, A. S.; Plückthun, High-Resolution Crystal Structure of Parathyroid Hormone 1 Receptor in Complex with a Peptide Agonist. Nature Structural and Molecular Biology 2018, 25 (12), 1086–1092. https://doi.org/10.1038/s41594-018-0151-4.

(15) Binkowski, B. F.; Butler, B. L.; Stecha, P. F.; Eggers, C. T.; Otto, P.; Zimmerman, K.; Vidugiris, G.; Wood, M. G.; Encell, L. P.; Fan, F.; Wood, K. v. A Luminescent Biosensor with Increased Dynamic Range for Intracellular CAMP. ACS Chemical Biology 2011, 6 (11), 1193–1197. https://doi.org/10.1021/cb200248h.

(16) Salahpour, A.; Espinoza, S.; Masri, B.; Lam, V.; Barak, L. S.; Gainetdinov, R. R. BRET Biosensors to Study GPCR Biology, Pharmacology, and Signal Transduction. Frontiers in Endocrinology 2012, 3 (105), 1–9. https://doi.org/10.3389/fendo.2012.00105.

(17) Gales, C.; Rebois, R. V.; Hogue, M.; Trieu, P.; Breit, A.; Hebert, T. E.; Bouvier, M. Real-Time Monitoring of Receptor and G-Protein Interactions in Living Cells. 2005, 2 (3), 177–184. https://doi.org/10.1038/NMETH743.

(18) Hamdan, F. F.; Audet, M.; Garneau, P.; Pelletier, J.; Bouvier, M. High-Throughput Screening of G Protein-Coupled Receptor Antagonists Using a Bioluminescence Resonance Energy Transfer 1-Based β-Arrestin2 Recruitment Assay. Journal of Biomolecular Screening 2005, 10 (5), 463–475. https://doi.org/10.1177/1087057105275344.

(19) Lohse, M. J.; Nuber, S.; Hoffmann, C. Fluorescence / Bioluminescence Resonance Energy Transfer Techniques to Study G-Protein-Coupled. Pharmacological Reviews 2012, 64 (2), 299–336. https://doi.org/10.1124/pr.110.004309.

(20) Kauk, M.; Hoffmann, C. Intramolecular and Intermolecular FRET Sensors for GPCRs –Monitoring Conformational Changes and Beyond. Trends in Pharmacological Sciences 2018, 39 (2), 123–135. https://doi.org/10.1016/j.tips.2017.10.011.

(21) Gardella, T. J.; Luck, M. D.; Wilson, A. K.; Keutmann, H. T.; Nussbaum, S. R.; Potts, J. T.; Kronenberg, H. M. Parathyroid Hormone (PTH)-PTH-Related Peptide Hybrid Peptides Reveal Functional Interactions between the 1-14 and 15-34 Domains of the Ligand. Journal of Biological Chemistry 1995, 270 (12), 6584–6588. https://doi.org/10.1074/jbc.270.12.6584.

(22) Gardella, T. J.; Luck, M. D.; Jensen, G. S.; Usdin, T. B.; Jüppner, H. Converting Parathyroid Hormone-Related Peptide (PTHrP) into a Potent PTH-2 Receptor Agonist. Journal of Biological Chemistry 1996, 271 (33), 19888–19893. https://doi.org/10.1074/jbc.271.33.19888.

(23) Dean, T.; Vilardaga, J. P.; Potts, J. T.; Gardella, T. J. Altered Selectivity of Parathyroid Hormone (PTH) and PTH-Related Protein (PTHrP) for Distinct Conformations of the PTH/PTHrP Receptor. Molecular Endocrinology 2008, 22 (1), 156–166. https://doi.org/10.1210/me.2007-0274.

(24) Hattersley, G.; Dean, T.; Corbin, B. A.; Bahar, H.; Gardella, T. J. Binding Selectivity of Abaloparatide for PTH-Type-1-Receptor Conformations and Effects on Downstream Signaling. Endocrinology 2016, 157 (1), 141–149. https://doi.org/10.1210/en.2015-1726.

(25) Dean, T.; Linglart, A.; Mahon, M. J.; Bastepe, M.; Jüppner, H.; Potts, J. T.; Gardella, T. J. Mechanisms of Ligand Binding to the Parathyroid Hormone (PTH)/PTH-Related Protein Receptor: Selectivity of a Modified PTH(1-15) Radioligand for GαS-Coupled Receptor Conformations. Molecular Endocrinology 2006, 20 (4), 931–943. https://doi.org/10.1210/me.2005-0349.

(26) Noda, H.; Okazaki, M.; Joyashiki, E.; Tamura, T.; Kawabe, Y.; Khatri, A.; Jueppner, H.; Potts, J. T.; Gardella, T. J.; Shimizu, M. Optimization of PTH / PTHrP Hybrid Peptides to Derive a Long-Acting PTH Analog (LA-PTH). JBMR Plus 2020, 4 (7), 1–10. https://doi.org/10.1002/jbm4.10367.

(27) Shimizu, N.; Guo, J.; Gardella, T. J. Parathyroid Hormone (PTH)-(1-14) and -(1-11) Analogs Conformationally Constrained by α-Aminoisobutyric Acid Mediate Full Agonist Responses via the Juxtamembrane Region of the PTH-1 Receptor. Journal of Biological Chemistry 2001, 276 (52), 49003–49012. https://doi.org/10.1074/jbc.M106827200.

(28) Grätz, L.; Tropmann, K.; Bresinsky, M.; Müller, C.; Bernhardt, G.; Pockes, S. NanoBRET Binding Assay for Histamine H2 Receptor Ligands Using Live Recombinant HEK293T Cells. Scientific Reports 2020, 10 (1), 1–10. https://doi.org/10.1038/s41598-020-70332-3.

(29) Bouzo-Lorenzo, M.; Stoddart, L. A.; Xia, L.; IJzerman, A. P.; Heitman, L. H.; Briddon, S. J.; Hill, S. J. A Live Cell NanoBRET Binding Assay Allows the Study of Ligand-Binding Kinetics to the Adenosine A3 Receptor. Purinergic Signalling 2019, 15 (2), 139–153. https://doi.org/10.1007/s11302-019-09650-9.

(30) Ricarte, F. R.; le Henaff, C.; Kolupaeva, V. G.; Gardella, T. J.; Partridge, N. C. Parathyroid Hormone(1–34) and Its Analogs Differentially Modulate Osteoblastic Rankl Expression via PKA/SIK2/SIK3 and PP1/PP2A–CRTC3 Signaling. Journal of Biological Chemistry 2018, 293 (52), 20200–20213. https://doi.org/10.1074/jbc.RA118.004751.

(31) Ferrandon, S.; Feinstein, T. N.; Castro, M.; Wang, B.; Bouley, R.; Potts, J. T.; Gardella, T. J.; Vilardaga, J. P. Sustained Cyclic AMP Production by Parathyroid Hormone Receptor Endocytosis. Nature Chemical Biology 2009, 5 (10), 734–742. https://doi.org/10.1038/nchembio.206.

(32) Garton, M.; Nim, S.; Stone, T. A.; Wang, K. E.; Deber, C. M.; Kim, P. M. Method to Generate Highly Stable D-Amino Acid Analogs of Bioactive Helical Peptides Using a Mirror Image of the Entire PDB. Proc Natl Acad Sci U S A 2018, 115 (7), 1505–1510. https://doi.org/10.1073/pnas.1711837115.

(33) Hilliker, S.; Wergedal, J. E.; Gruber, H. E.; Bettica, P.; Baylink, D. J. Truncation of the Amino Terminus of PTH Alters Its Anabolic Activity on Bone in Vivo. Bone 1996, 19 (5), 469–477. https://doi.org/10.1016/S8756-3282(96)00230-X.

(34) Sutkeviciute, I.; Clark, L. J.; White, A. D.; Gardella, T. J.; Vilardaga, J. P. PTH/PTHrP Receptor Signaling, Allostery, and Structures. Trends in Endocrinology and Metabolism 2019, 30 (11), 860–874. https://doi.org/10.1016/j.tem.2019.07.011.

(35) Hu, Y. B.; Dammer, E. B.; Ren, R. J.; Wang, G. The Endosomal-Lysosomal System: From Acidification and Cargo Sorting to Neurodegeneration. Translational Neurodegeneration 2015, 4 (1), 1–10. https://doi.org/10.1186/s40035-015-0041-1.

(36) Pioszak, A. A.; Xu, H. E. Molecular Recognition of Parathyroid Hormone by Its G Protein-Coupled Receptor. Proc Natl Acad Sci U S A 2008, 105 (13), 5034–5039. https://doi.org/10.1073/pnas.0801027105.

(37) Pioszak, A. A.; Parker, N. R.; Gardella, T. J.; Xu, H. E. Structural Basis for Parathyroid Hormone-Related Protein Binding to the Parathyroid Hormone Receptor and Design of Conformation-Selective Peptides. Journal of Biological Chemistry 2009, 284 (41), 28382–28391. https://doi.org/10.1074/jbc.M109.022905.

(38) Vilardaga, J.-P.; Nikolaev, V. O.; Lohse, M. J.; Palm, D.; Castro, M. Turn-on Switch in Parathyroid Hormone Receptor by a Two-Step Parathyroid Hormone Binding Mechanism. Proceedings of the National Academy of Sciences 2005, 102 (44), 16084–16089. https://doi.org/10.1073/pnas.0503942102.

(39) Kuo, S. W.; Rimando, M. G.; Liu, Y. S.; Lee, O. K. Intermittent Administration of Parathyroid Hormone 1-34 Enhances Osteogenesis of Human Mesenchymal Stem Cells by Regulating Protein Kinase Cδ. International Journal of Molecular Sciences 2017, 18 (10), 1–16. https://doi.org/10.3390/ijms18102221.

(40) Nakajima, A.; Shimoji, N.; Shiomi, K.; Shimizu, S.; Moriya, H.; Einhorn, T. A.; Yamazaki, M. Mechanisms for the Enhancement of Fracture Healing in Rats Treated with Intermittent Low-Dose Human Parathyroid Hormone (1-34). Journal of Bone and Mineral Research 2002, 17 (11), 2038–2047. https://doi.org/10.1359/jbmr.2002.17.11.2038.

(41) Cary, B. P.; Deganutti, G.; Zhao, P.; Truong, T. T.; Piper, S. J.; Liu, X.; Belousoff, M. J.; Danev, R.; Sexton, P. M.; Wootten, D.; Gellman, S. H. Structural and Functional Diversity among Agonist-Bound States of the GLP-1 Receptor. Nature Chemical Biology 2022, 18 (3), 256–263. https://doi.org/10.1038/s41589-021-00945-w.

