## Supplementary Information for "Kinetic and Thermodynamic Insights on Agonist Interactions with the Parathyroid Hormone Receptor-1 from a New NanoBRET assay"

**Peptide Characterization**

PTH[1-34]-N33K^TMR^ ; SVSEIQLMHN LGKHLNSMER VEWLRKKLQD VH-K^TMR^-F-NH_2_

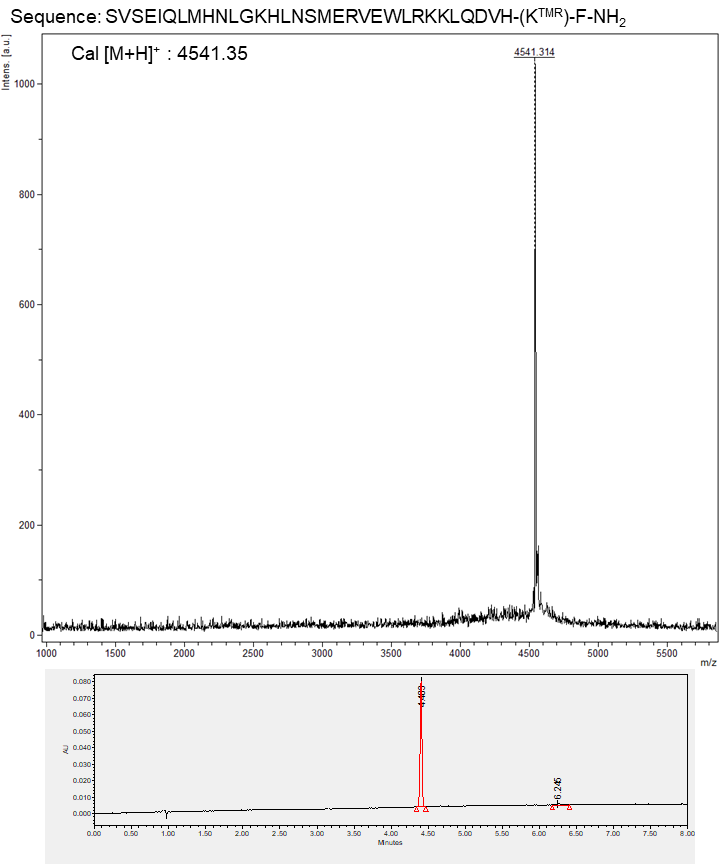

MALDI-TOF-MS (top) and UPLC (bottom) analysis of the PTH[1-34]-N33K^TMR^ peptide. UPLC chromatogram was obtained with a ACQUITY UPLC BEH C18 column (2.1mm X 100mm) eluted with a linear gradient of 10-90% acetonitrile in water (0.1% TFA) applied over 8 min at a flow rate of 0.3 mL/min. Purity >95%.

PTH[1-34]-35K^TMR^ ; SVSEIQLMHN LGKHLNSMER VEWLRKKLQD VHNF-K^TMR^-NH_2_

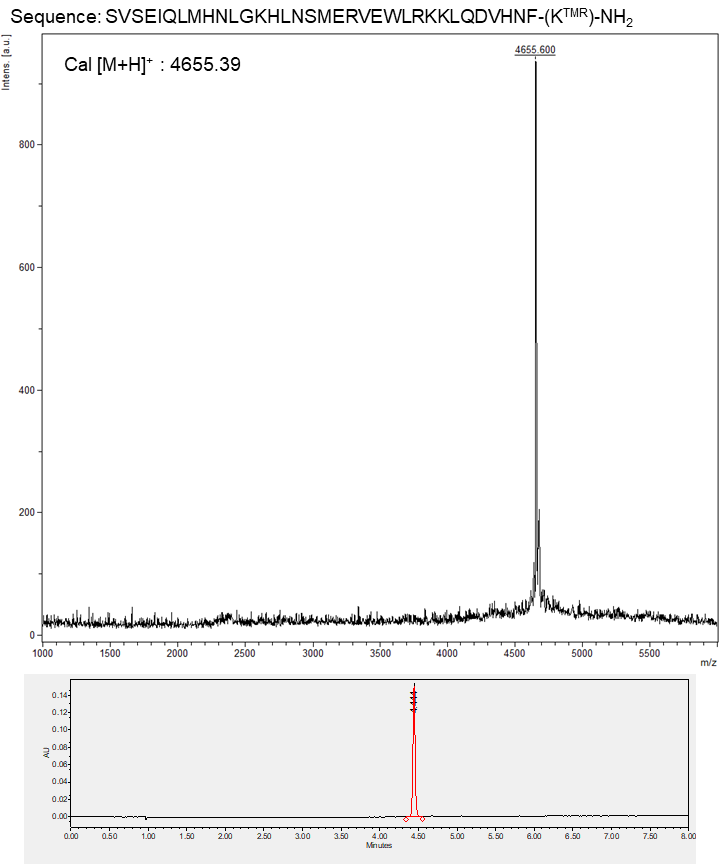

MALDI-TOF-MS (top) and UPLC (bottom) analysis of the PTH[1-34]-35K^TMR^ peptide. UPLC chromatogram was obtained with a ACQUITY UPLC BEH C18 column (2.1mm X 100mm) eluted with a linear gradient of 10-90% acetonitrile in water (0.1% TFA) applied over 8 min at a flow rate of 0.3 mL/min. Purity >95%.

PTHrP[1-12]-PTH[13-34]-35K^TMR^ ; AVSEHQLLHD KGKHLNSMER VEWLRKKLQD VHNF-K^TMR^-NH_2_

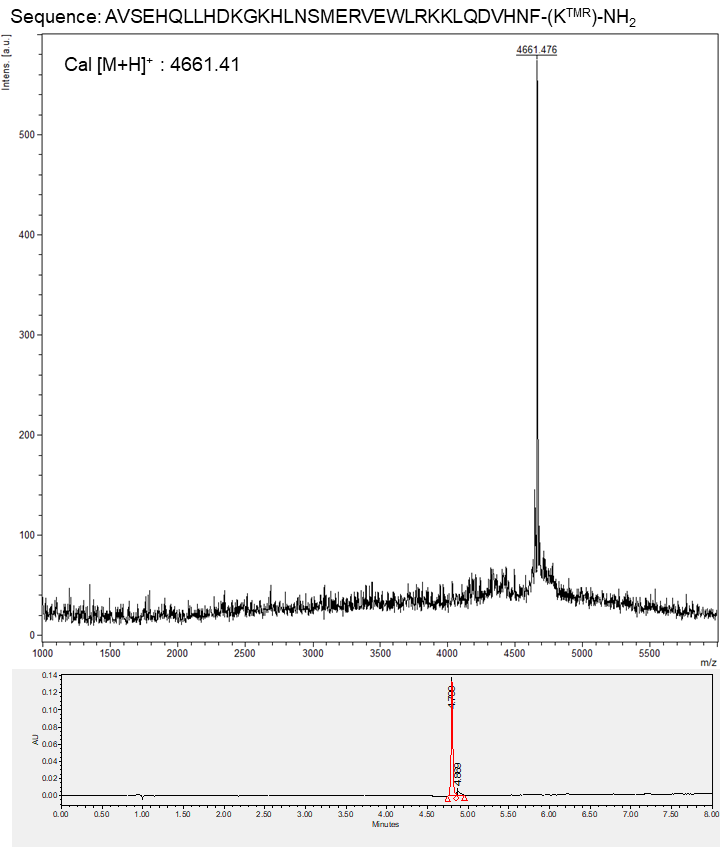

MALDI-TOF-MS (top) and UPLC (bottom) analysis of the PTHrP[1-12]-PTH[13-34]-35K^TMR^ peptide. UPLC chromatogram was obtained with a ACQUITY UPLC BEH C18 column (2.1mm X 100mm) eluted with a linear gradient of 10-90% acetonitrile in water (0.1% TFA) applied over 8 min at a flow rate of 0.3 mL/min. Purity >95%.

PTHrP[1-14]-PTH[15-34]-35K^TMR^ ; AVSEHQLLHD KGKSLNSMER VEWLRKKLQD VHNF-K^TMR^-NH_2_

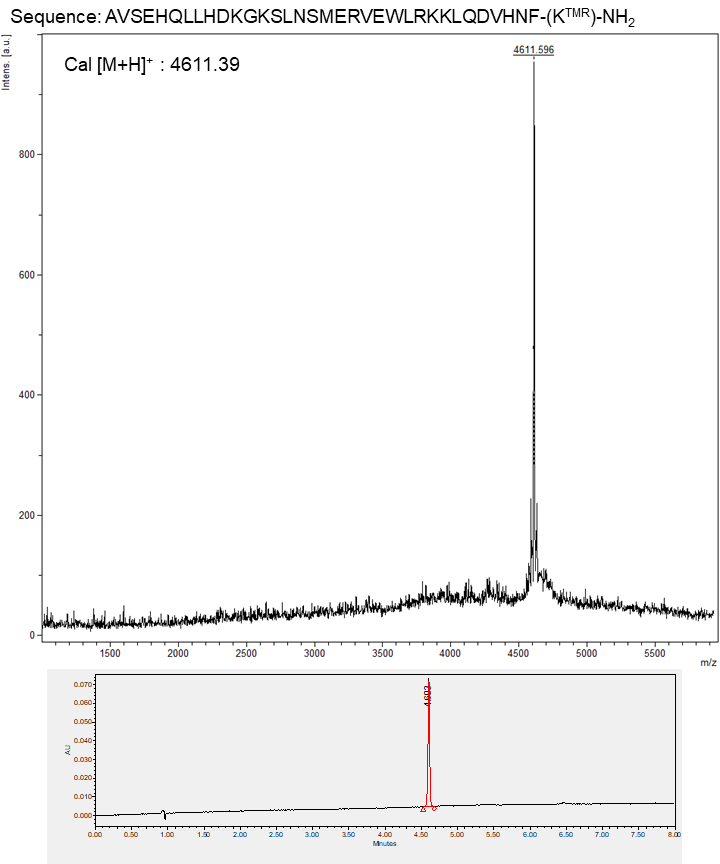

MALDI-TOF-MS (top) and UPLC (bottom) analysis of the PTHrP[1-14]-PTH[15-34]-35K^TMR^ peptide. UPLC chromatogram was obtained with a ACQUITY UPLC BEH C18 column (2.1mm X 100mm) eluted with a linear gradient of 10-90% acetonitrile in water (0.1% TFA) applied over 8 min at a flow rate of 0.3 mL/min. Purity >95%.

PTHrP[1-16]-PTH[17-34]-35K^TMR^ ; AVSEHQLLHD KGKSIQSMER VEWLRKKLQD VHNF-K^TMR^-NH_2_

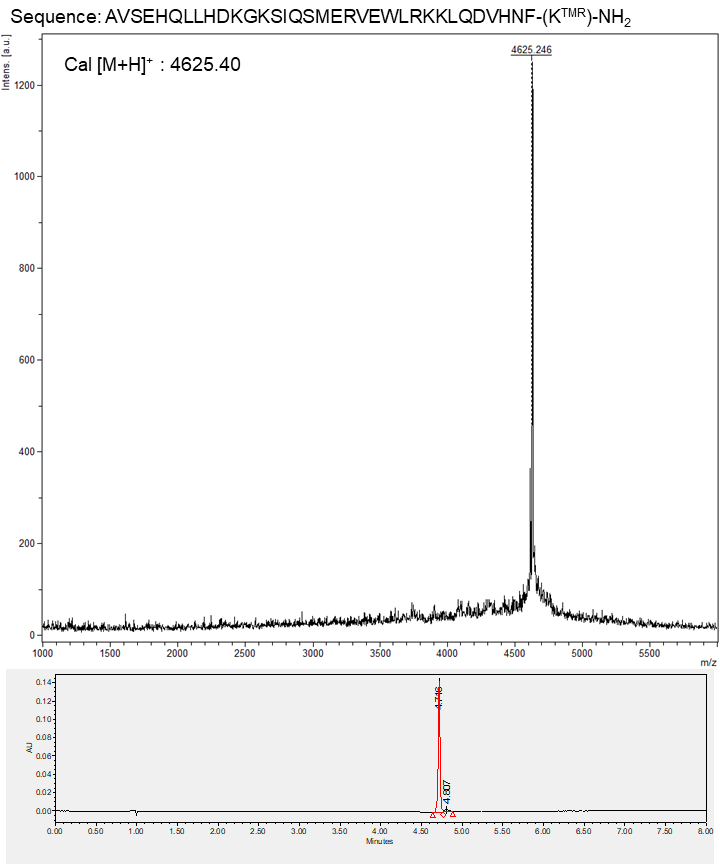
 MALDI-TOF-MS (top) and UPLC (bottom) analysis of the PTHrP[1-16]-PTH[17-34]-35K^TMR^ peptide. UPLC chromatogram was obtained with a ACQUITY UPLC BEH C18 column (2.1mm X 100mm) eluted with a linear gradient of 10-90% acetonitrile in water (0.1% TFA) applied over 8 min at a flow rate of 0.3 mL/min. Purity >95%.

PTHrP[1-18]-PTH[19-34]-35K^TMR^ ; AVSEHQLLHD KGKSIQDLER VEWLRKKLQD VHNF-K^TMR^-NH_2_

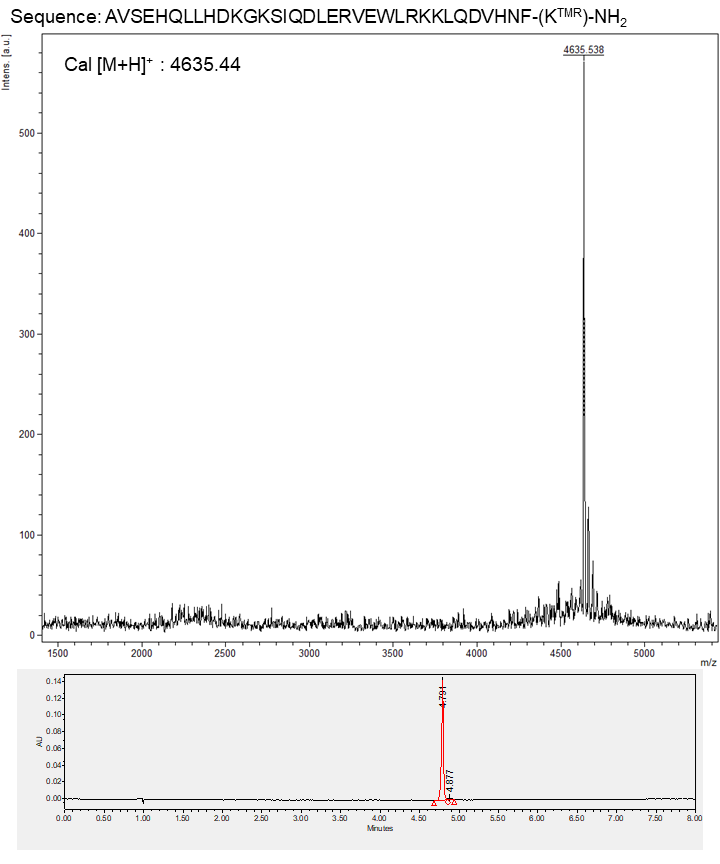

MALDI-TOF-MS (top) and UPLC (bottom) analysis of the PTHrP[1-18]-PTH[19-34]-35K^TMR^ peptide. UPLC chromatogram was obtained with a ACQUITY UPLC BEH C18 column (2.1mm X 100mm) eluted with a linear gradient of 10-90% acetonitrile in water (0.1% TFA) applied over 8 min at a flow rate of 0.3 mL/min. Purity >95%.

PTH[1-12]-PTHrP[13-36]-E35K^TMR^ ; SVSEIQLMHN LGKSIQDLRR RFFLHHLIAE IHTA-K^TMR^-I-NH_2_

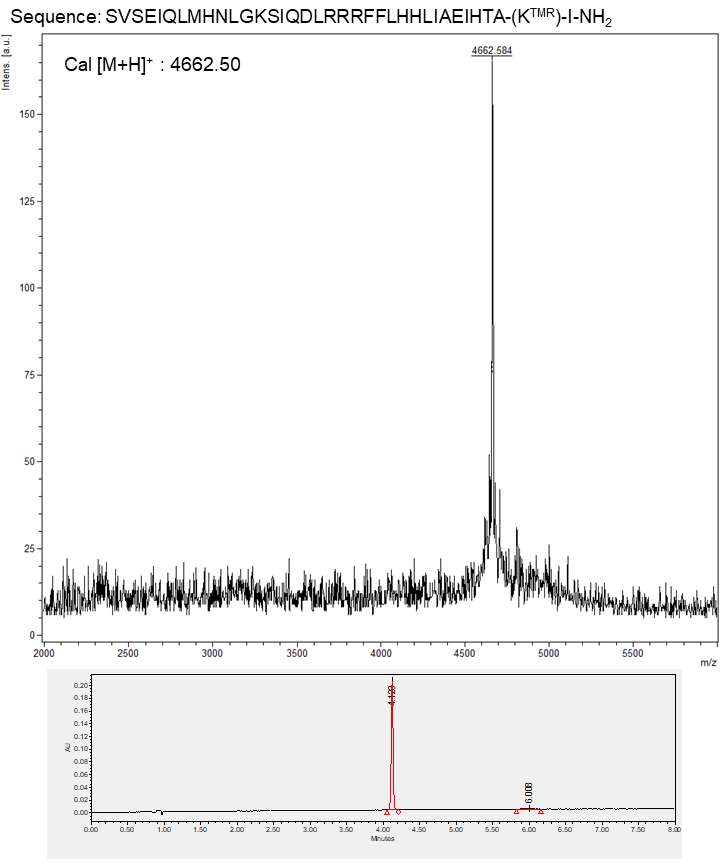

MALDI-TOF-MS (top) and UPLC (bottom) analysis of the PTH[1-12]-PTHrP[13-36]-E35K^TMR^ peptide. UPLC chromatogram was obtained with a ACQUITY UPLC BEH C18 column (2.1mm X 100mm) eluted with a linear gradient of 10-90% acetonitrile in water (0.1% TFA) applied over 8 min at a flow rate of 0.3 mL/min. Purity >95%.

PTH[1-14]-PTHrP[15-36]-E35K^TMR^ ; SVSEIQLMHN LGKHIQDLRR RFFLHHLIAE IHTA-K^TMR^-I-NH_2_

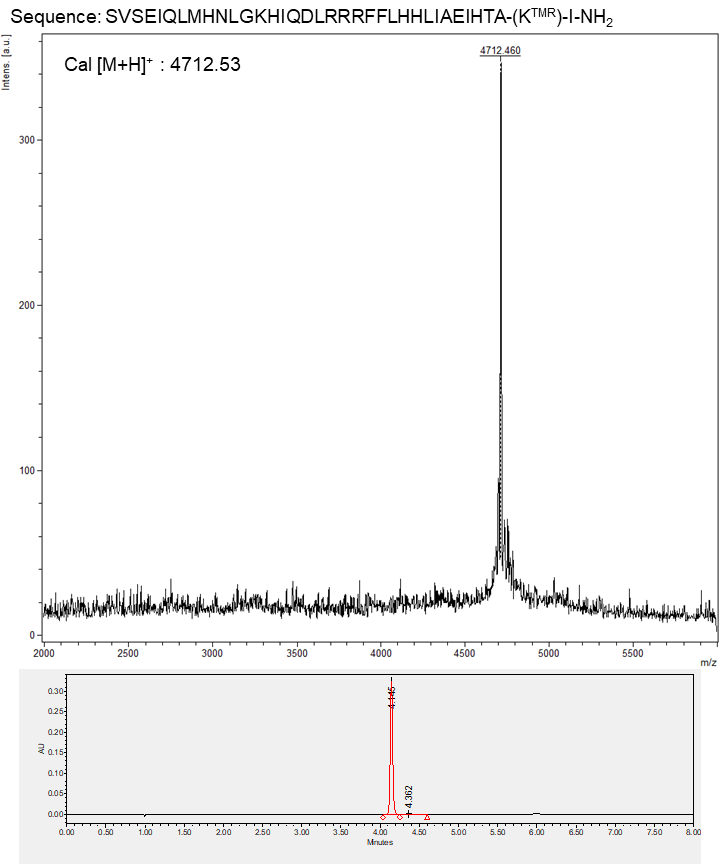

MALDI-TOF-MS (top) and UPLC (bottom) analysis of the PTH[1-14]-PTHrP[15-36]-E35K^TMR^ peptide. UPLC chromatogram was obtained with a ACQUITY UPLC BEH C18 column (2.1mm X 100mm) eluted with a linear gradient of 10-90% acetonitrile in water (0.1% TFA) applied over 8 min at a flow rate of 0.3 mL/min. Purity >95%.

PTH[1-16]-PTHrP[17-36]-E35K^TMR^ ; SVSEIQLMHN LGKHLNDLRR RFFLHHLIAE IHTA-K^TMR^-I-NH_2_

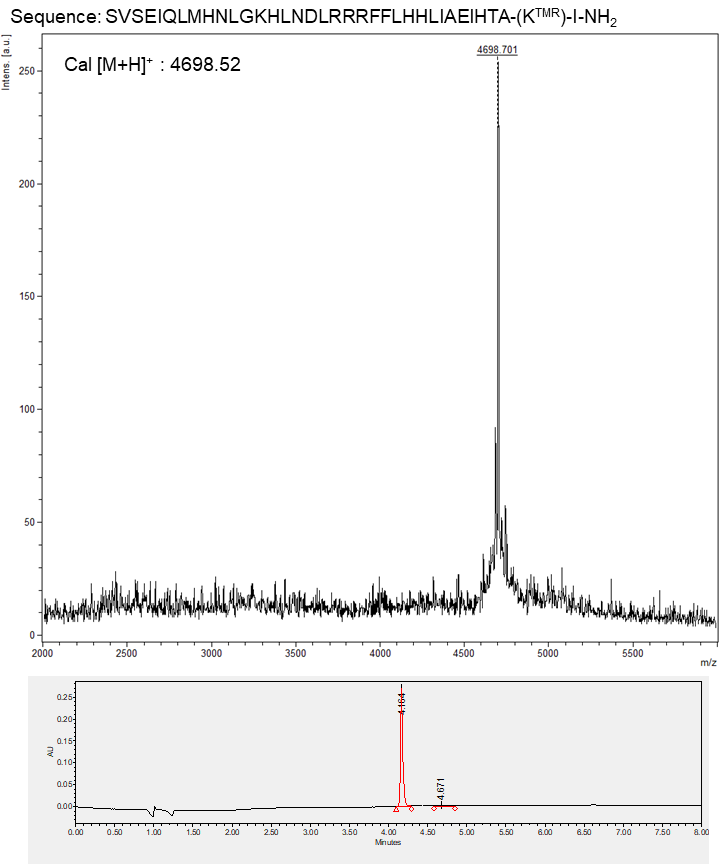

MALDI-TOF-MS (top) and UPLC (bottom) analysis of the PTH[1-16]-PTHrP[17-36]-E35K^TMR^ peptide. UPLC chromatogram was obtained with a ACQUITY UPLC BEH C18 column (2.1mm X 100mm) eluted with a linear gradient of 10-90% acetonitrile in water (0.1% TFA) applied over 8 min at a flow rate of 0.3 mL/min. Purity >95%.

PTH[1-18]-PTHrP[19-36]-E35K^TMR^ ; SVSEIQLMHN LGKHLNSMRR RFFLHHLIAE IHTA-K^TMR^-I-NH_2_

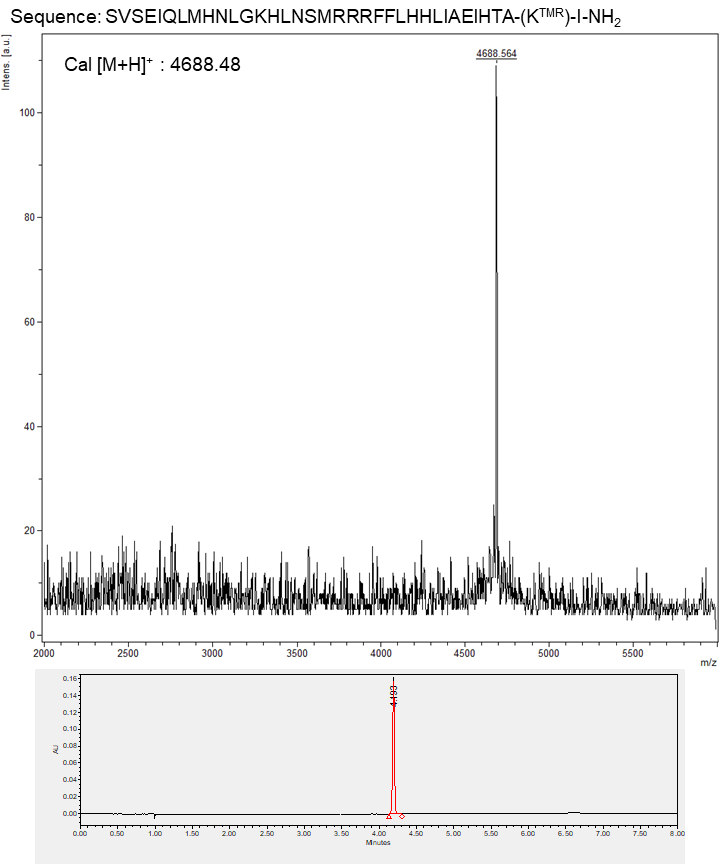

MALDI-TOF-MS (top) and UPLC (bottom) analysis of the PTH[1-18]-PTHrP[19-36]-E35K^TMR^ peptide. UPLC chromatogram was obtained with a ACQUITY UPLC BEH C18 column (2.1mm X 100mm) eluted with a linear gradient of 10-90% acetonitrile in water (0.1% TFA) applied over 8 min at a flow rate of 0.3 mL/min. Purity >95%.

PTHrP[1-34]-35K^TMR^ ; AVSEHQLLHD KGKSIQDLRR RFFLHHLIAE IHTA-K^TMR^-NH_2_

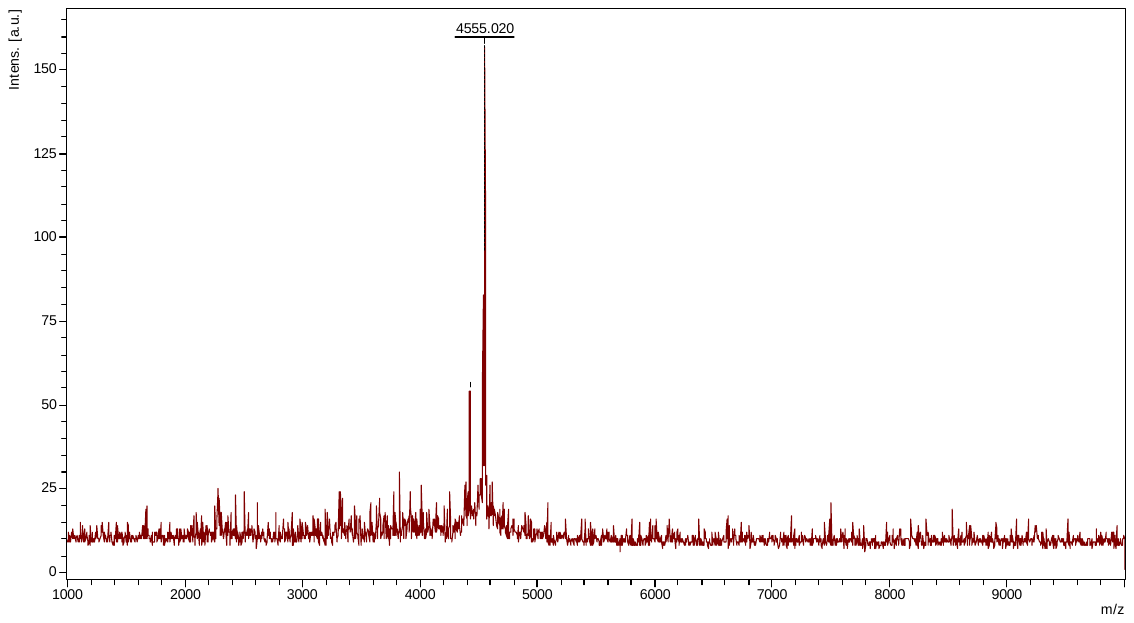

Cal [M+H]^+^ = 4555.29

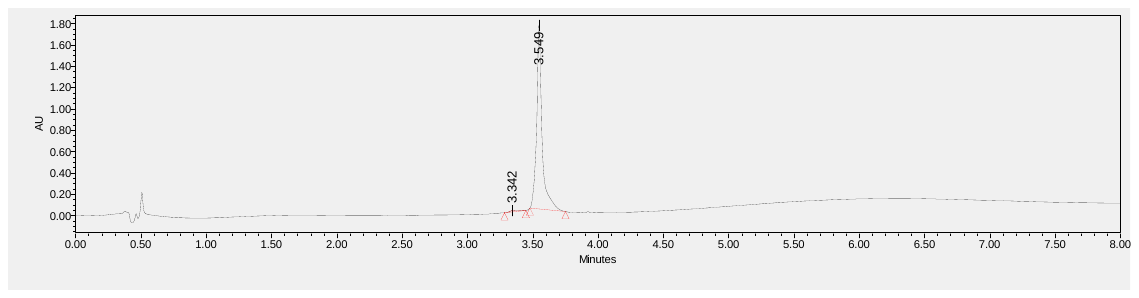

MALDI-TOF-MS (top) and UPLC (bottom) analysis of the PTHrP[1-34]-35K^TMR^ peptide. UPLC chromatogram was obtained with a ACQUITY UPLC BEH C18 column (2.1mm X 100mm) eluted with a linear gradient of 10-90% acetonitrile in water (0.1% TFA) applied over 8 min at a flow rate of 0.3 mL/min. Purity >95%.

LA-PTH[1-34]-35K^TMR^ ; AVAEIQLMHQ RAKWIQDARR RAFLHKLIAE IHTA-K^TMR^-NH_2_

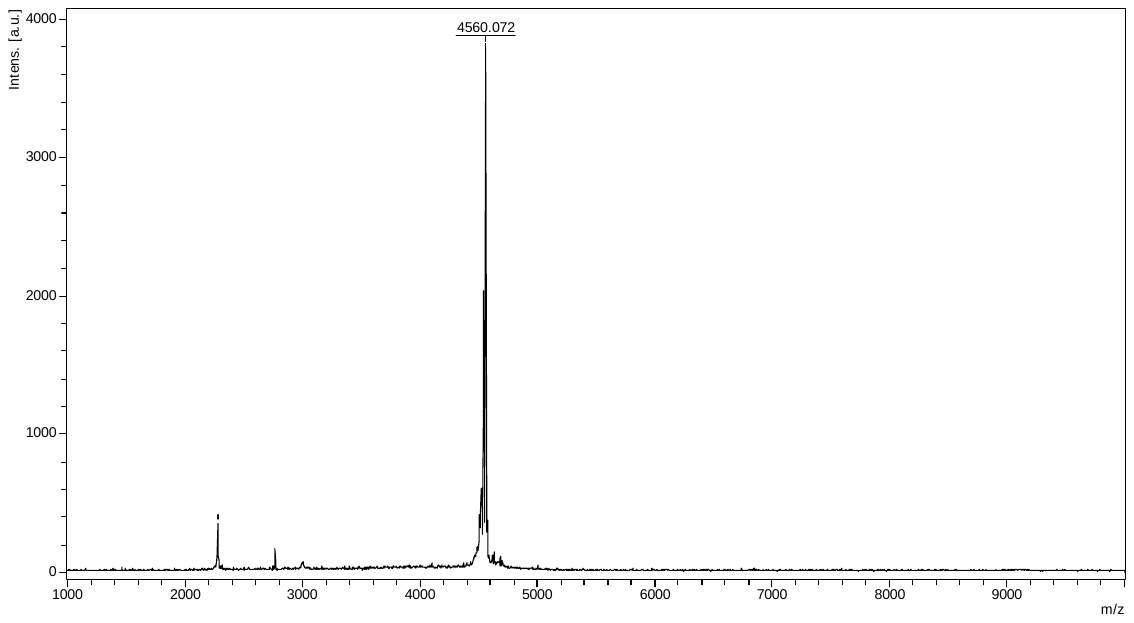

Cal [M+H]^+^ = 4560.34

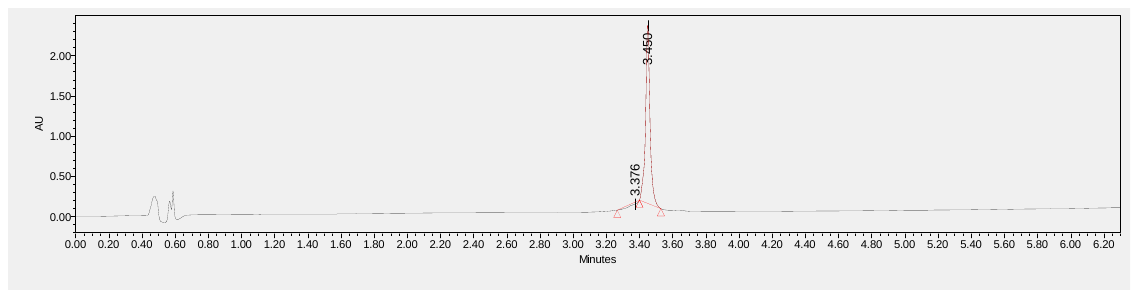

MALDI-TOF-MS (top) and UPLC (bottom) analysis of the LA-PTH[1-34]-35K^TMR^ peptide. UPLC chromatogram was obtained with a ACQUITY UPLC BEH C18 column (2.1mm X 100mm) eluted with a linear gradient of 10-90% acetonitrile in water (0.1% TFA) applied over 6 min at a flow rate of 0.3 mL/min. Purity >95%.

PTHrP[1-34]-13K^TMR^ ; AVSEHQLLHD KG-K^TMR^-SIQDLRR RFFLHHLIAE IHTA-NH_2_

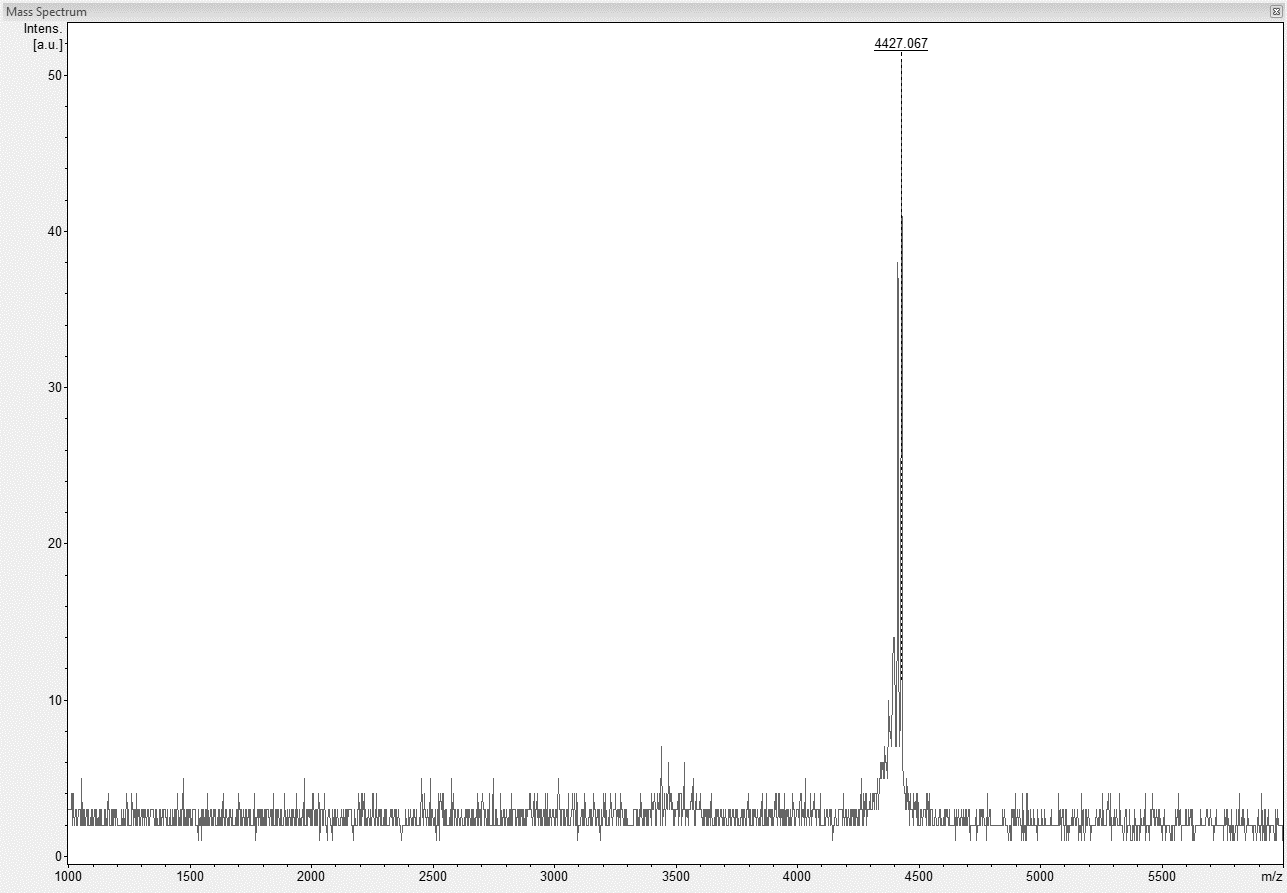

Cal [M+H]^+^ = 4427.19

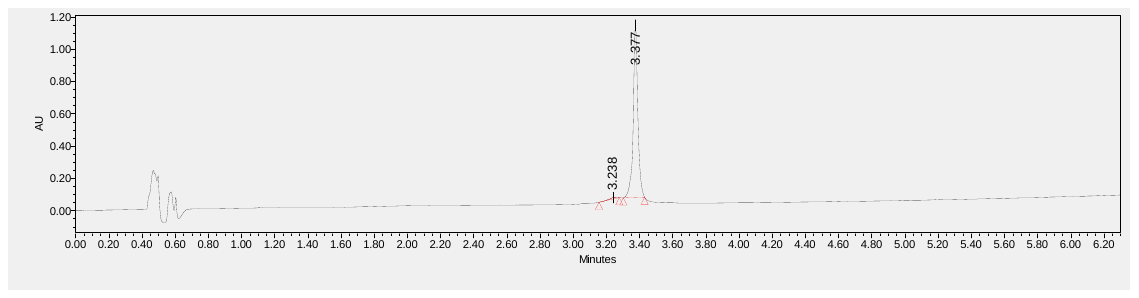

MALDI-TOF-MS (top) and UPLC (bottom) analysis of the PTHrP[1-34]-13K^TMR^ peptide. UPLC chromatogram was obtained with a ACQUITY UPLC BEH C18 column (2.1mm X 100mm) eluted with a linear gradient of 10-90% acetonitrile in water (0.1% TFA) applied over 6 min at a flow rate of 0.3 mL/min. Purity >95%.

PTH[2-34]-35K^TMR^ ; VSEIQLMHN LGKHLNSMER VEWLRKKLQD VHNF-K^TMR^-NH_2_

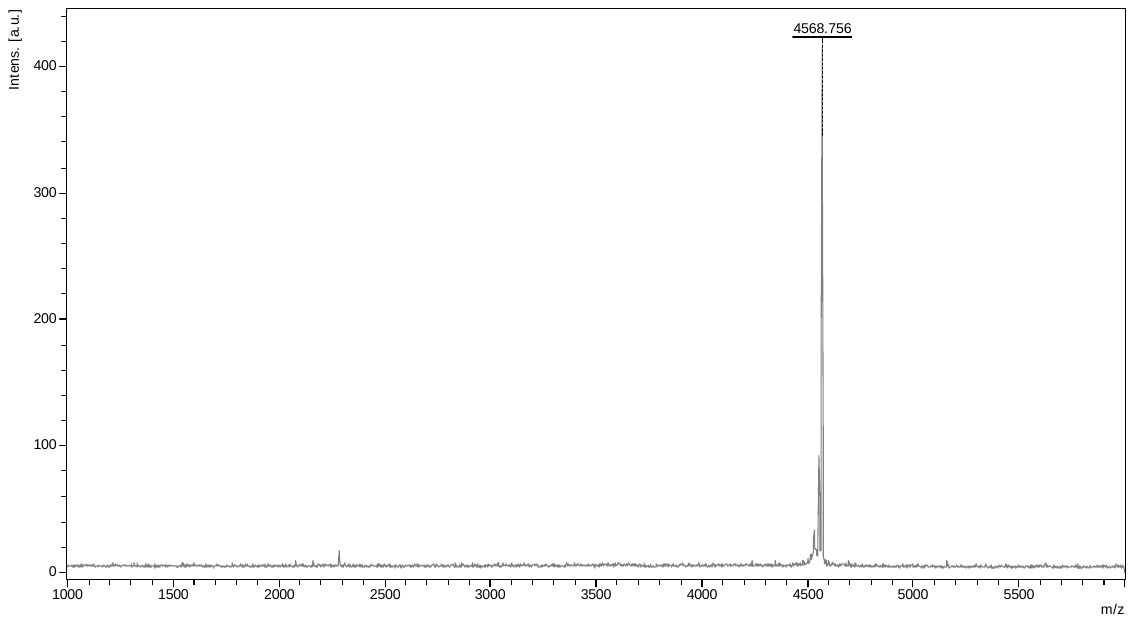

Cal [M+H]^+^ = 4568.2

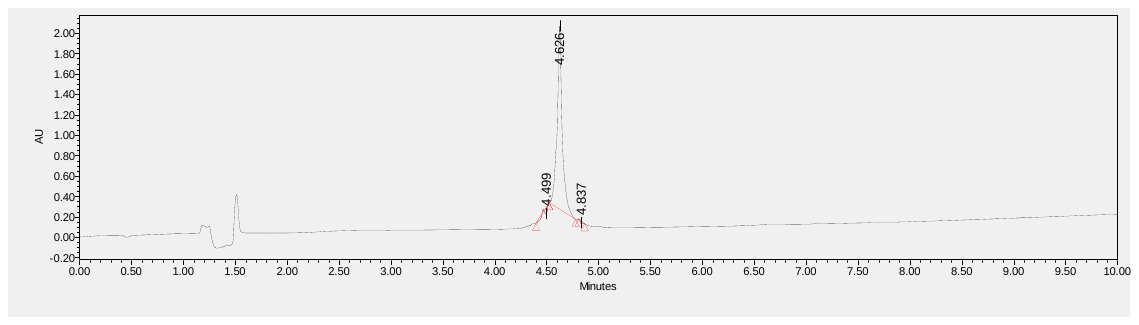

MALDI-TOF-MS (top) and UPLC (bottom) analysis of the PTH[2-34]-35K^TMR^ peptide. UPLC chromatogram was obtained with a ACQUITY UPLC BEH C18 column (2.1mm X 100mm) eluted with a linear gradient of 10-90% acetonitrile in water (0.1% TFA) applied over 8 min at a flow rate of 0.3 mL/min. Purity >95%.

PTH[3-34]-35K^TMR^ ; SEIQLMHN LGKHLNSMER VEWLRKKLQD VHNF-K^TMR^-NH_2_

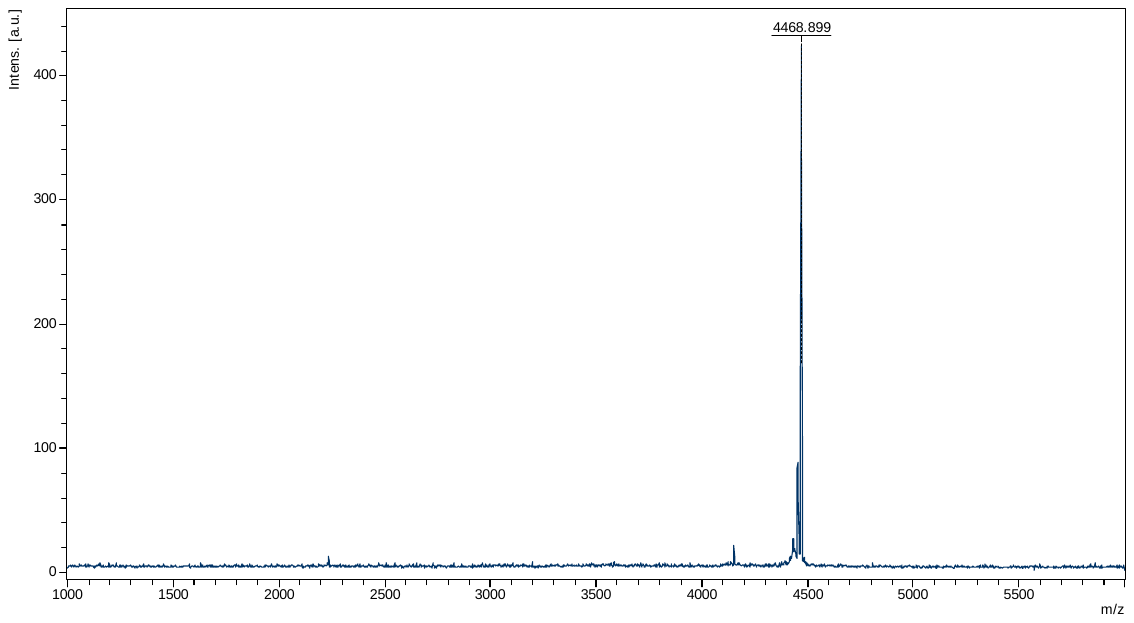

Cal [M+H]^+^ = 4468.15

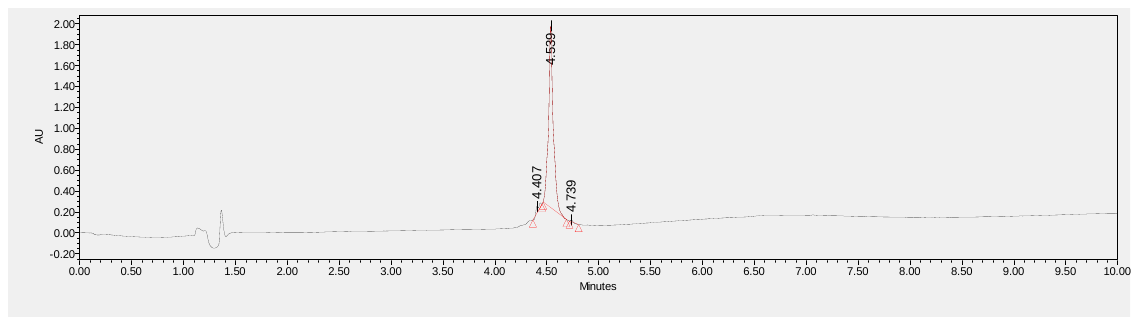

MALDI-TOF-MS (top) and UPLC (bottom) analysis of the PTH[3-34]-35K^TMR^ peptide. UPLC chromatogram was obtained with a ACQUITY UPLC BEH C18 column (2.1mm X 100mm) eluted with a linear gradient of 10-90% acetonitrile in water (0.1% TFA) applied over 8 min at a flow rate of 0.3 mL/min. Purity >95%.

M-PTH[1-14]-15K^TMR^ ; UVUEIQLMHQ R^h^AKW-K^TMR^-NH_2_ (U: aminoisobutyric acid, R^h^: homoarginine)

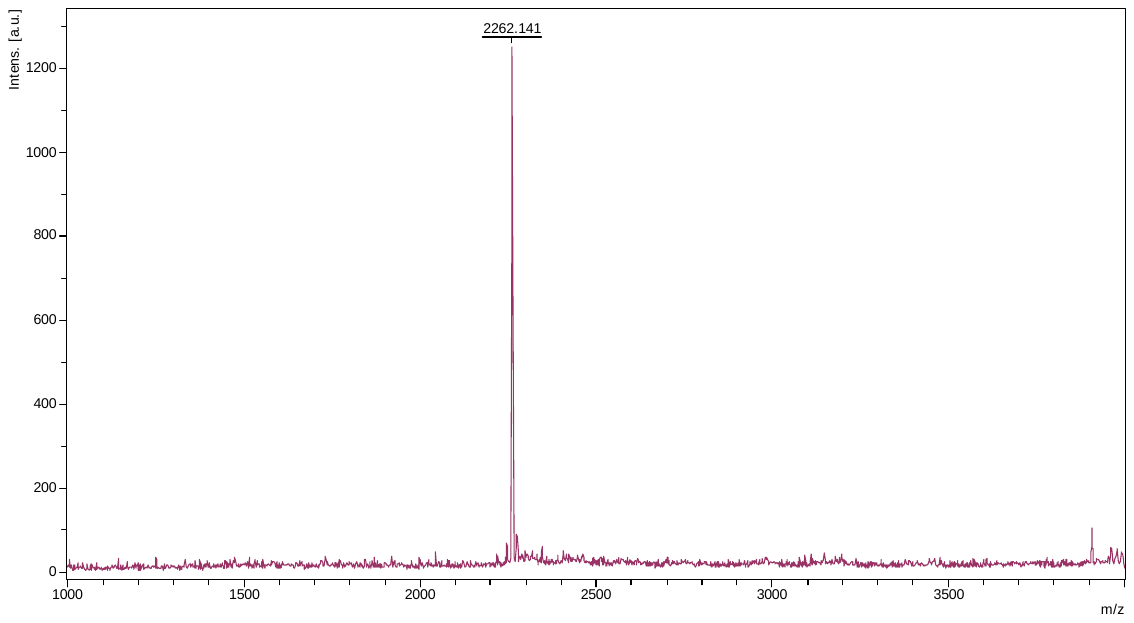

Cal [M+H]^+^ = 2262.04

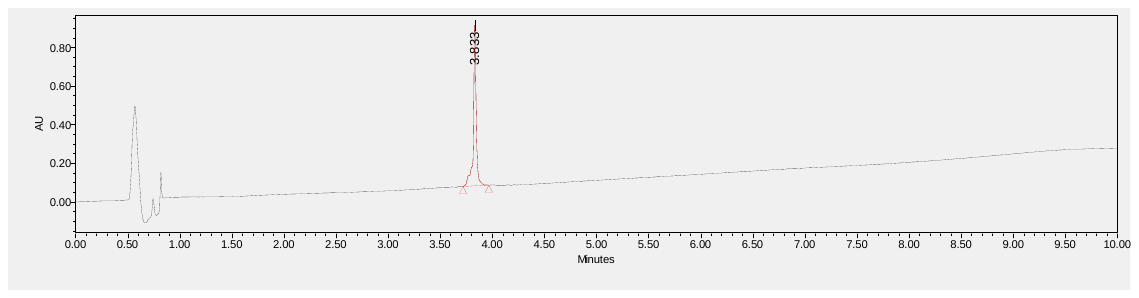

MALDI-TOF-MS (top) and UPLC (bottom) analysis of the M-PTH[1-14]-15K^TMR^ peptide. UPLC chromatogram was obtained with a ACQUITY UPLC BEH C18 column (2.1mm X 100mm) eluted with a linear gradient of 10-90% acetonitrile in water (0.1% TFA) applied over 8 min at a flow rate of 0.3 mL/min. Purity >95%.

PTH[1-34]I5H-35K^TMR^ ; SVSEHQLMHN LGKHLNSMER VEWLRKKLQD VHNF-K^TMR^-NH_2_

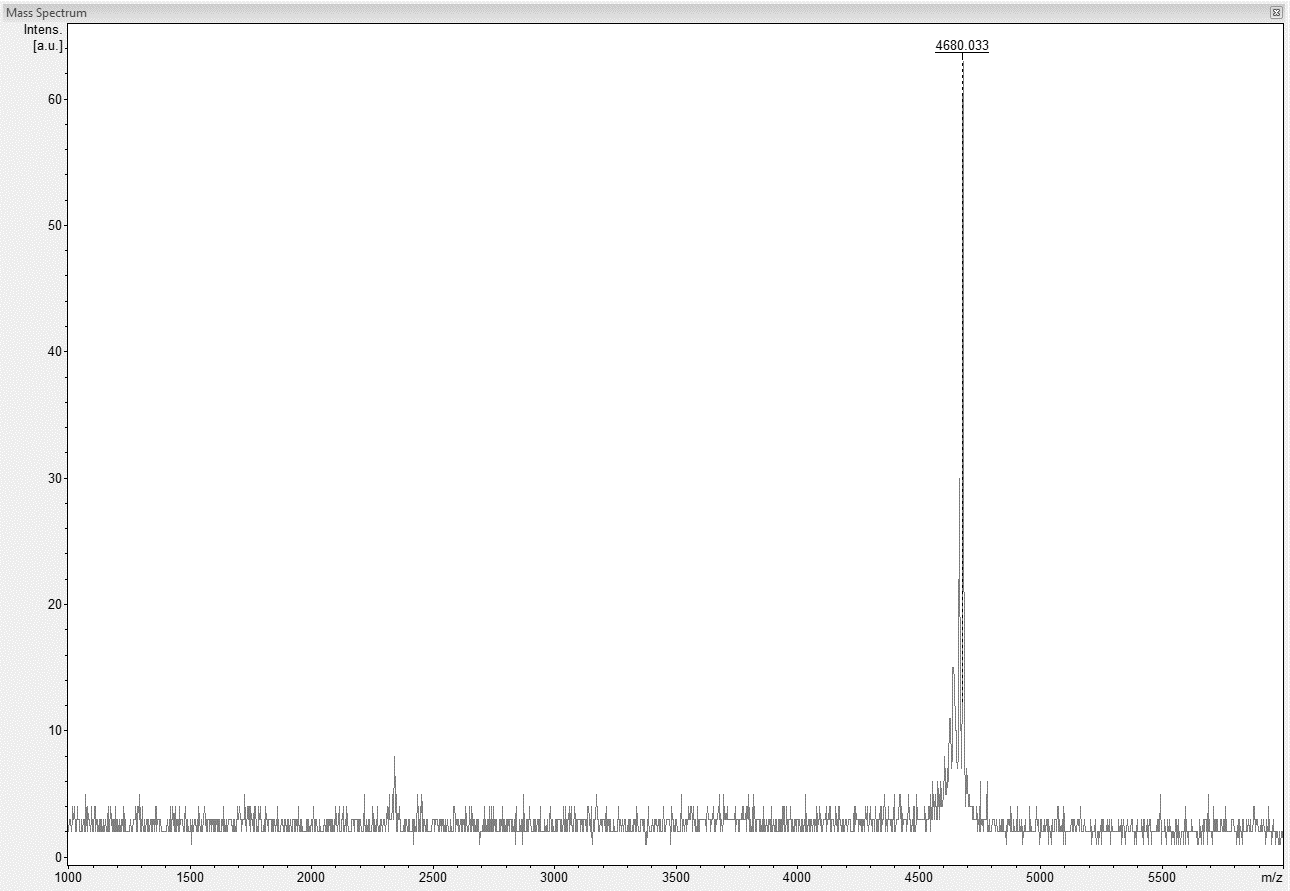

Cal [M+H]^+^ = 4679.22

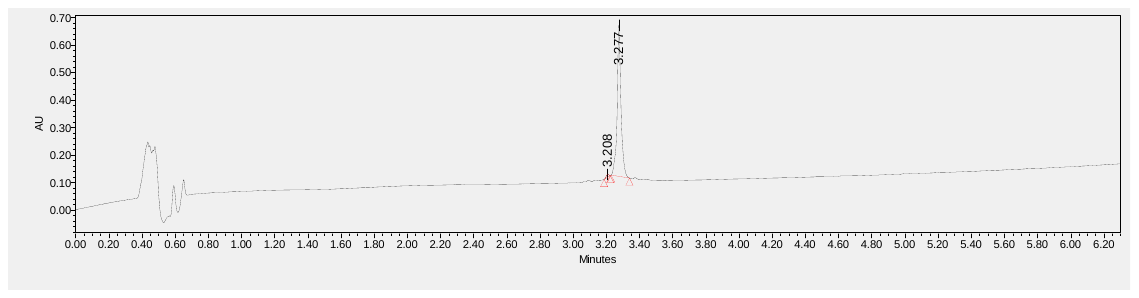

MALDI-TOF-MS (top) and UPLC (bottom) analysis of the PTH[1-34]I5H-35K^TMR^ peptide. UPLC chromatogram was obtained with a ACQUITY UPLC BEH C18 column (2.1mm X 100mm) eluted with a linear gradient of 10-90% acetonitrile in water (0.1% TFA) applied over 6 min at a flow rate of 0.3 mL/min. Purity >95%.

PTHrP[1-36]H5I-35K^TMR^ ; AVSEIQLLHD KGKSIQDLRR RFFLHHLIAE IHTA-K^TMR^-I-NH_2_

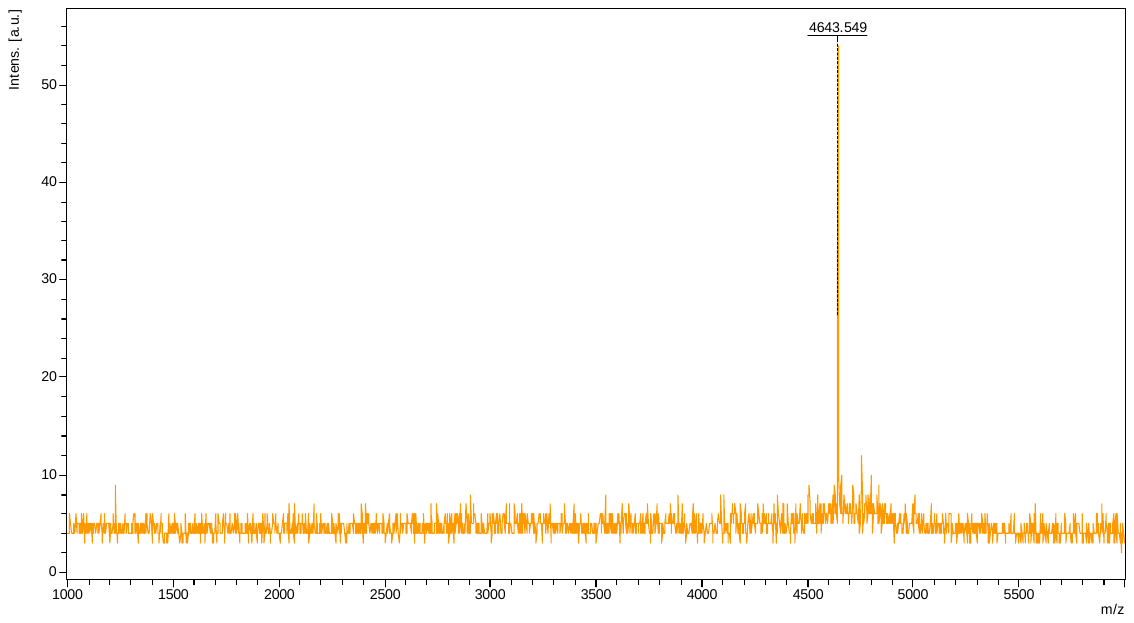

Cal [M+H]^+^ = 4643.40

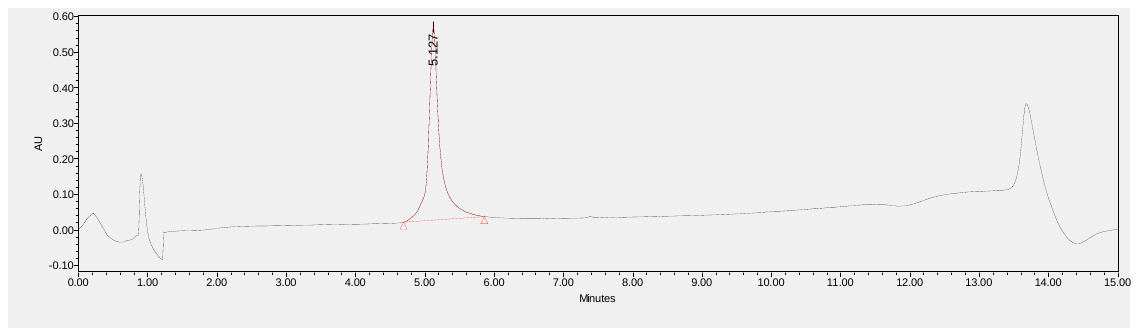

MALDI-TOF-MS (top) and UPLC (bottom) analysis of the PTHrP[1-36]H5I-35K^TMR^ peptide. UPLC chromatogram was obtained with a ACQUITY UPLC BEH C18 column (2.1mm X 100mm) eluted with a linear gradient of 10-90% acetonitrile in water (0.1% TFA) applied over 10 min at a flow rate of 0.3 mL/min. Purity >95%.

D-PTH ; aGlavadsvGmllmylpewrnhkGdlmreirsrkvs-NH_2_ (lowercase: D-configuration)

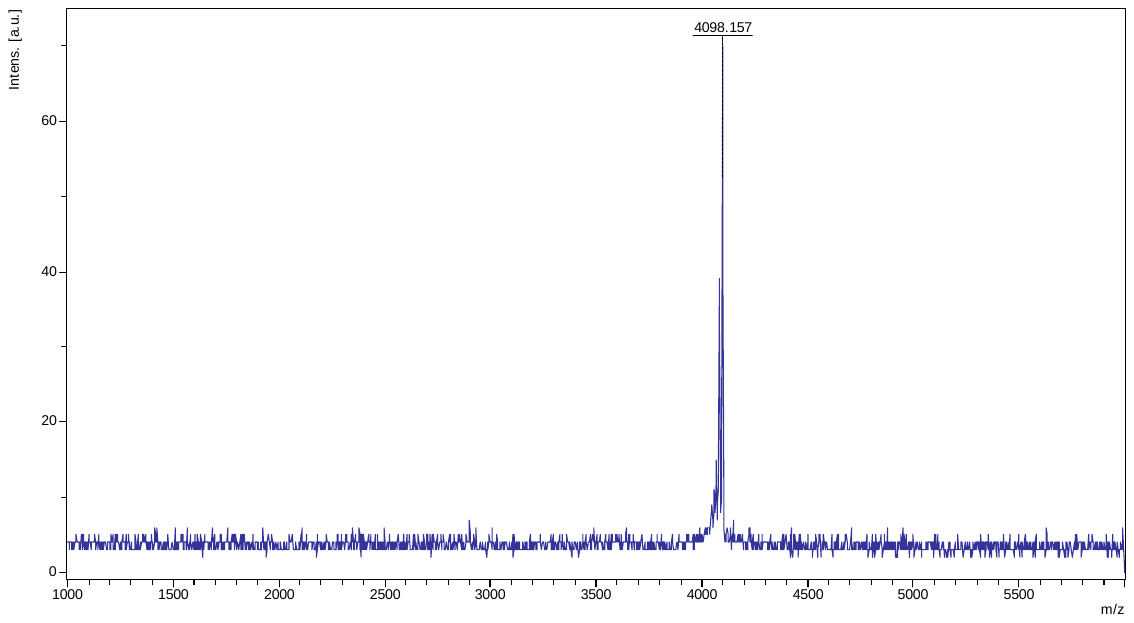

Cal [M+H]^+^ = 4098.17

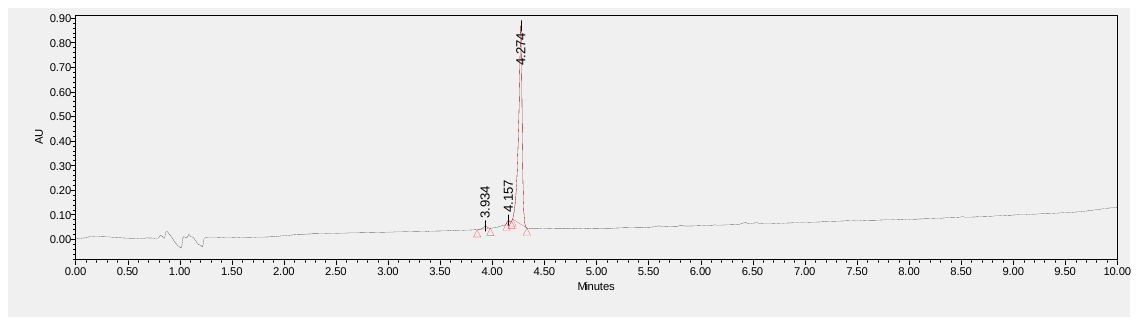

MALDI-TOF-MS (top) and UPLC (bottom) analysis of the D-PTH peptide. UPLC chromatogram was obtained with a ACQUITY UPLC BEH C18 column (2.1mm X 100mm) eluted with a linear gradient of 10-90% acetonitrile in water (0.1% TFA) applied over 8 min at a flow rate of 0.3 mL/min. Purity >95%.

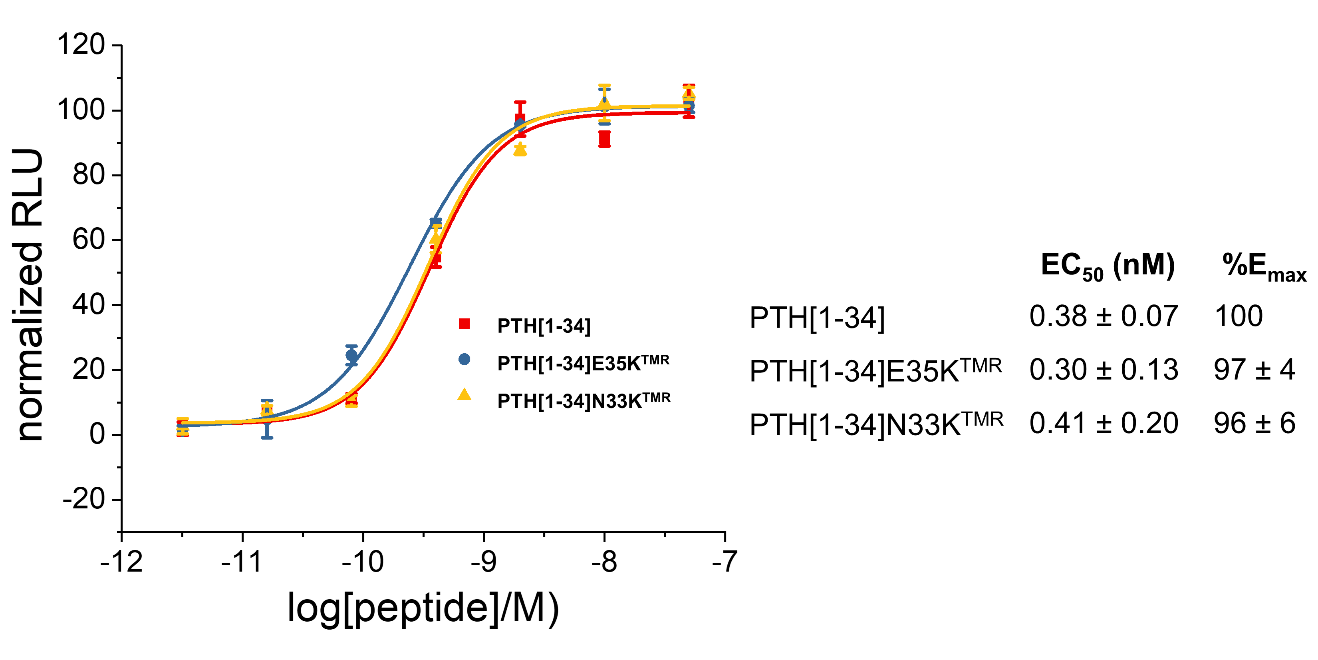

Figure S1. The cAMP formation stimulated by peptides were monitored by Glosensor in HEK293 cells. The luminescence response of vehicle control is normalized to 0%, while maximal response of unlabeled PTH(1-34) is normalized to 100%.

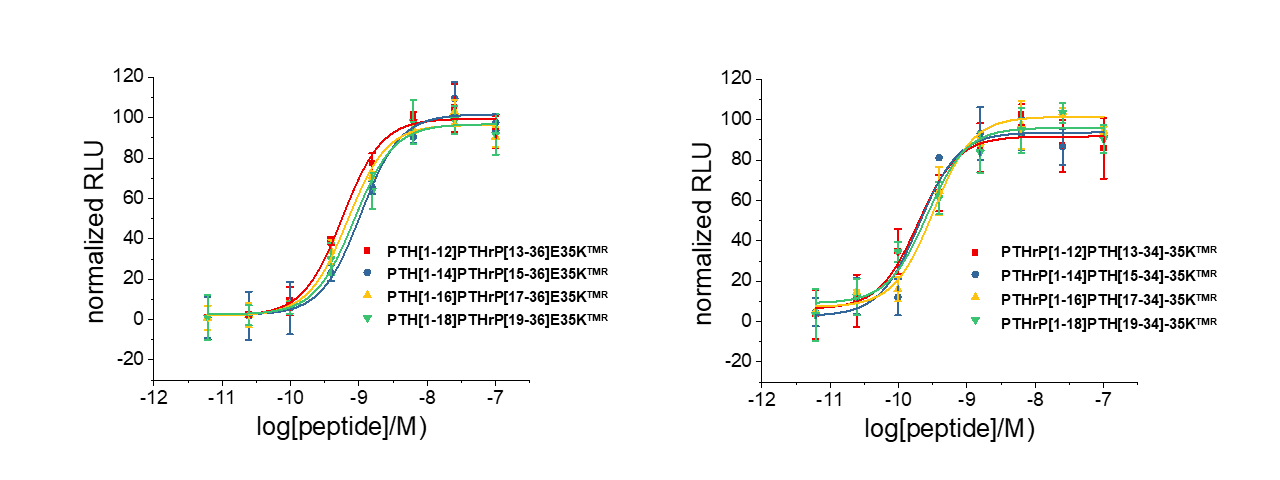

Figure S2. The cAMP formation stimulated by peptides was monitored by Glosensor in HEK293 cells. The luminescence response of vehicle control is normalized to 0%, while maximal response of unlabeled PTH(1-34) is normalized to 100%.

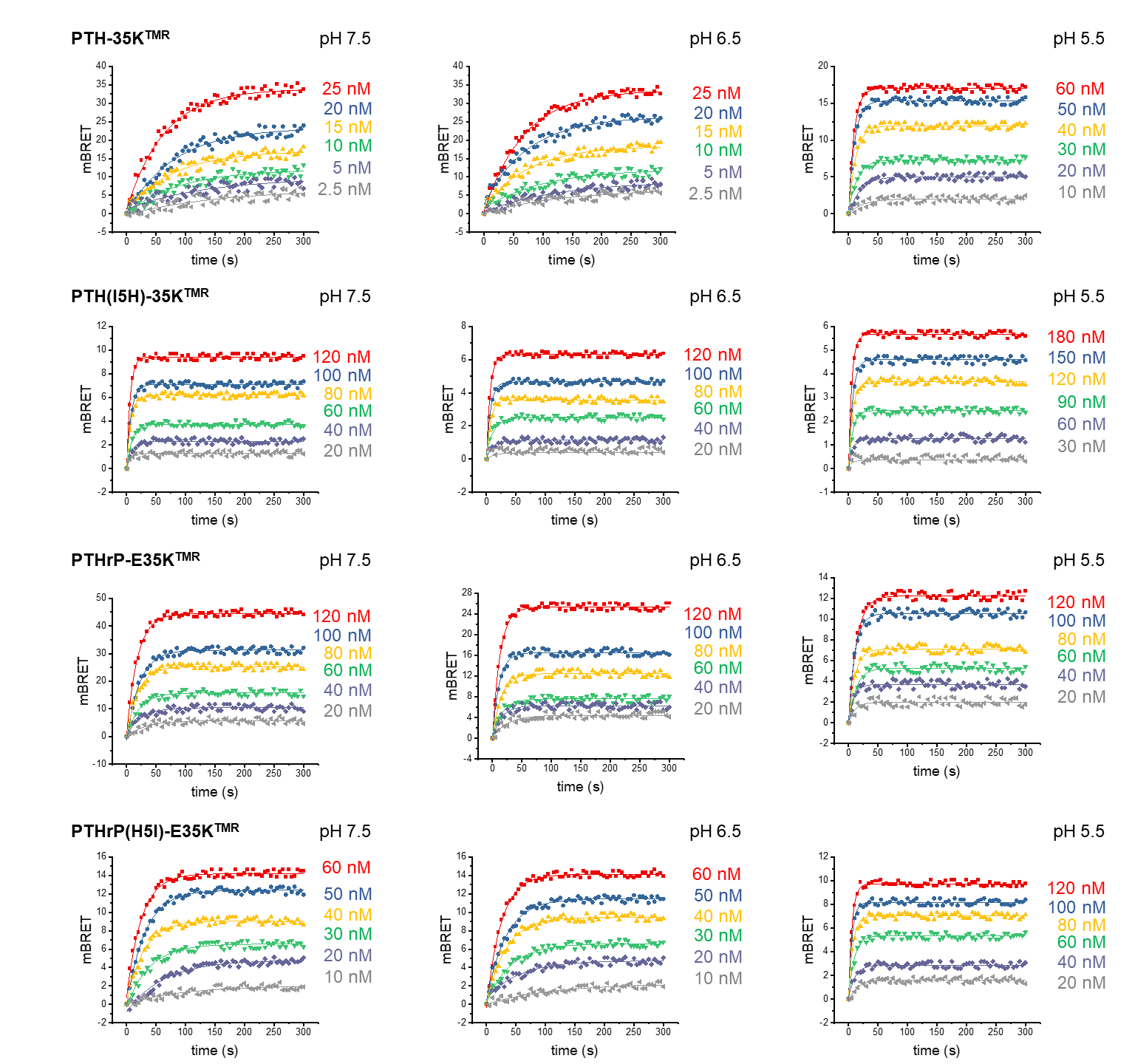

Figure S3. BRET curves showing the binding of the indicated peptides to nanoLuc-PTHR1 stably expressed on HEK293. Experiments were performed in 137 mM NaCl, 1 mM Ca^2+^, 0.5 mM Mg^2+^, and 0.1% (w/v) BSA buffered by 10 mM HEPES (pH 7.5), MES (pH 6.5) or acetate (pH 5.5).

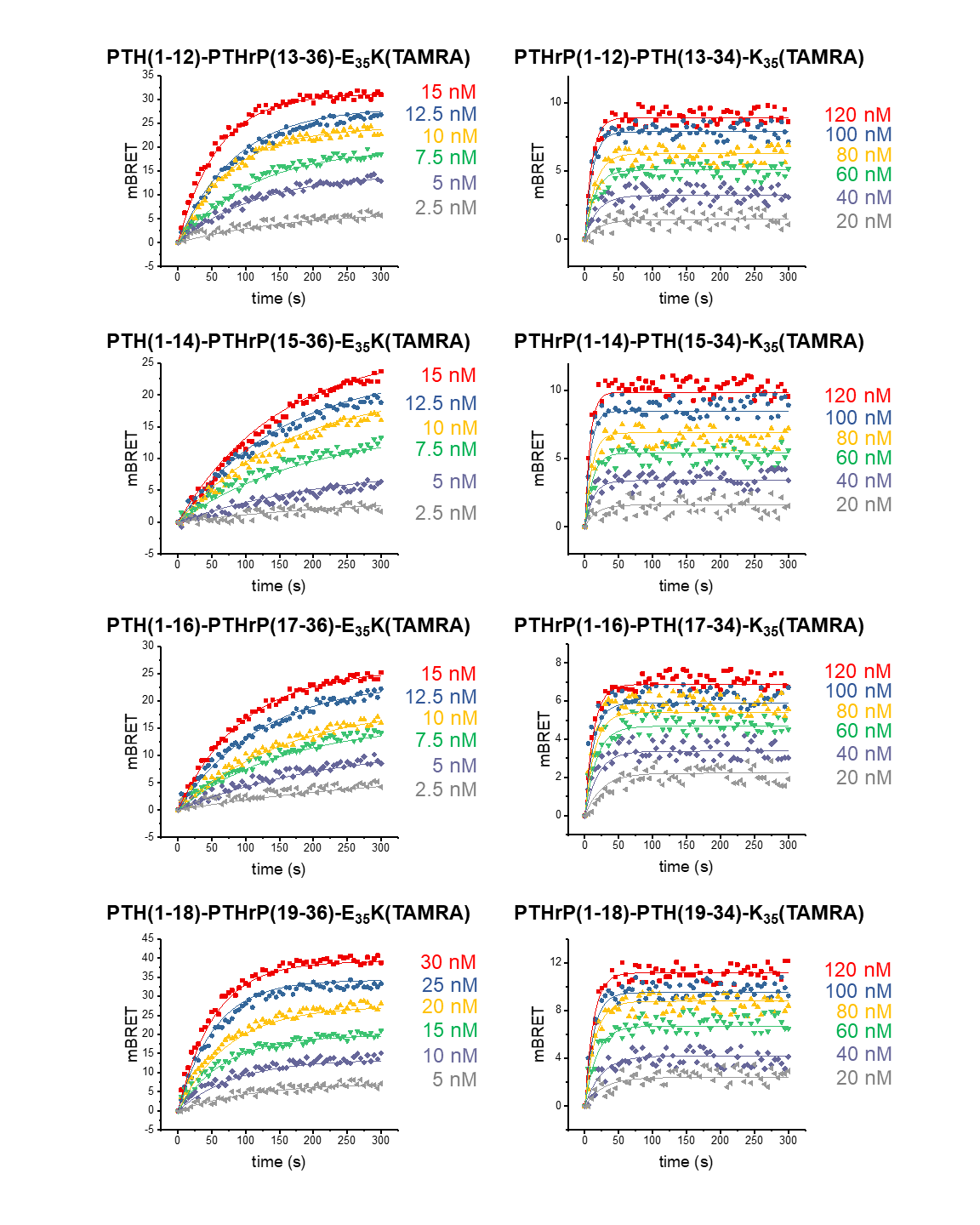

Figure S4. BRET curves showing the binding of the indicated hybrid peptides to nanoLuc-PTHR1 stably expressed on HEK293. Experiments were performed in 1 mM Ca^2+^, 0.5 mM Mg^2+^, 0.02% (w/v) NaN_3_, 0.1% (w/v) BSA with PBS.

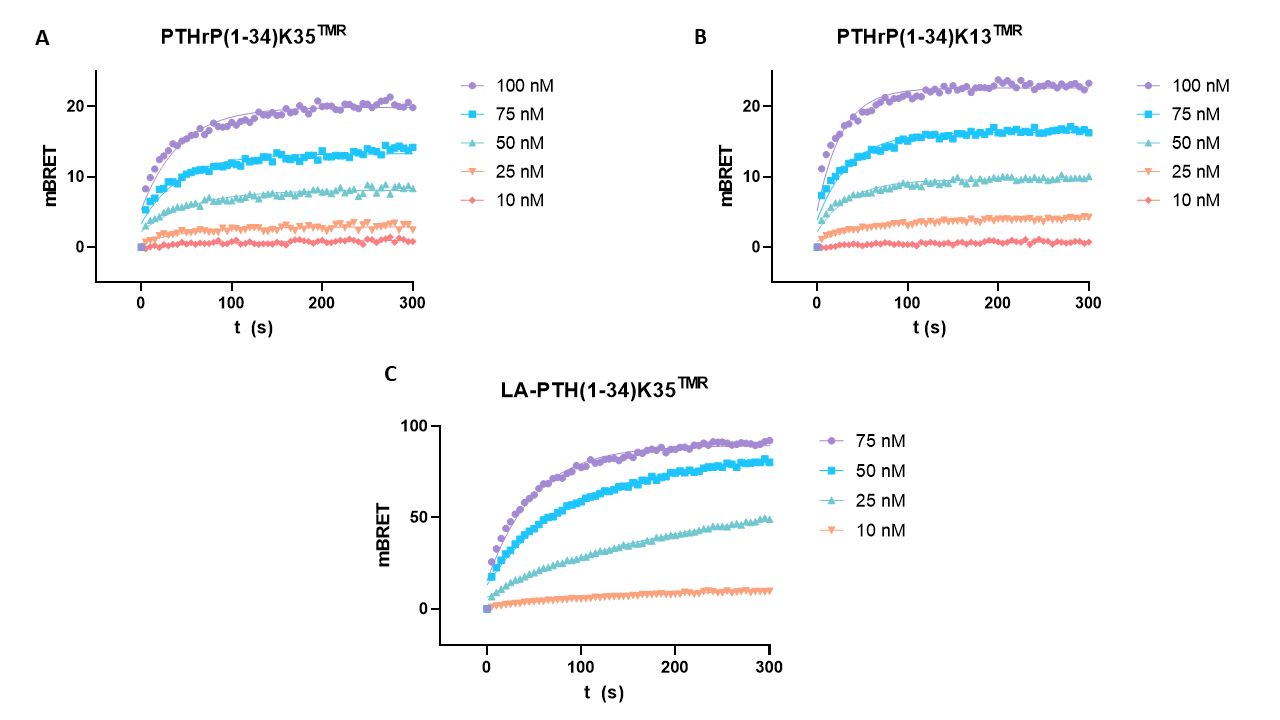

Figure S5. BRET curves showing the binding of the indicated peptides to nanoLuc-PTHR1 stably expressed on HEK293 cells. Experiments were performed in 1 mM Ca^2+^, 0.5 mM Mg^2+^, 0.02% (w/v) NaN_3_, 0.1% (w/v) BSA with PBS.

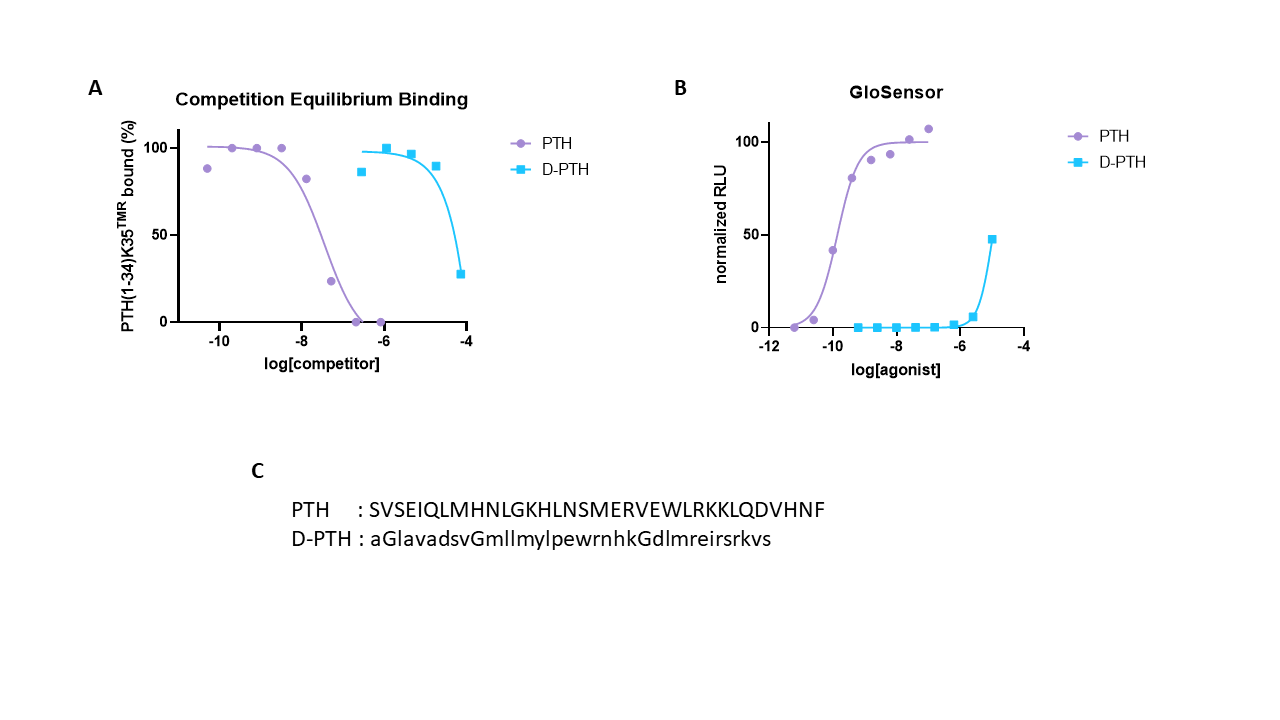

Figure S6. (A) Competition equilibrium binding of PTH(1-34) and D-PTH on nLuc-PTHR1 metabolically poisoned cells (0.02% NaN_3_ (w/v)). (B) cAMP measurement of PTH(1-34) and D-PTH on GP2.3 cells which stably express the GloSensor construct and hPTHR1. (C) Sequence of PTH(1-34) and D-PTH, where the lowercase letters represent D-configuration.

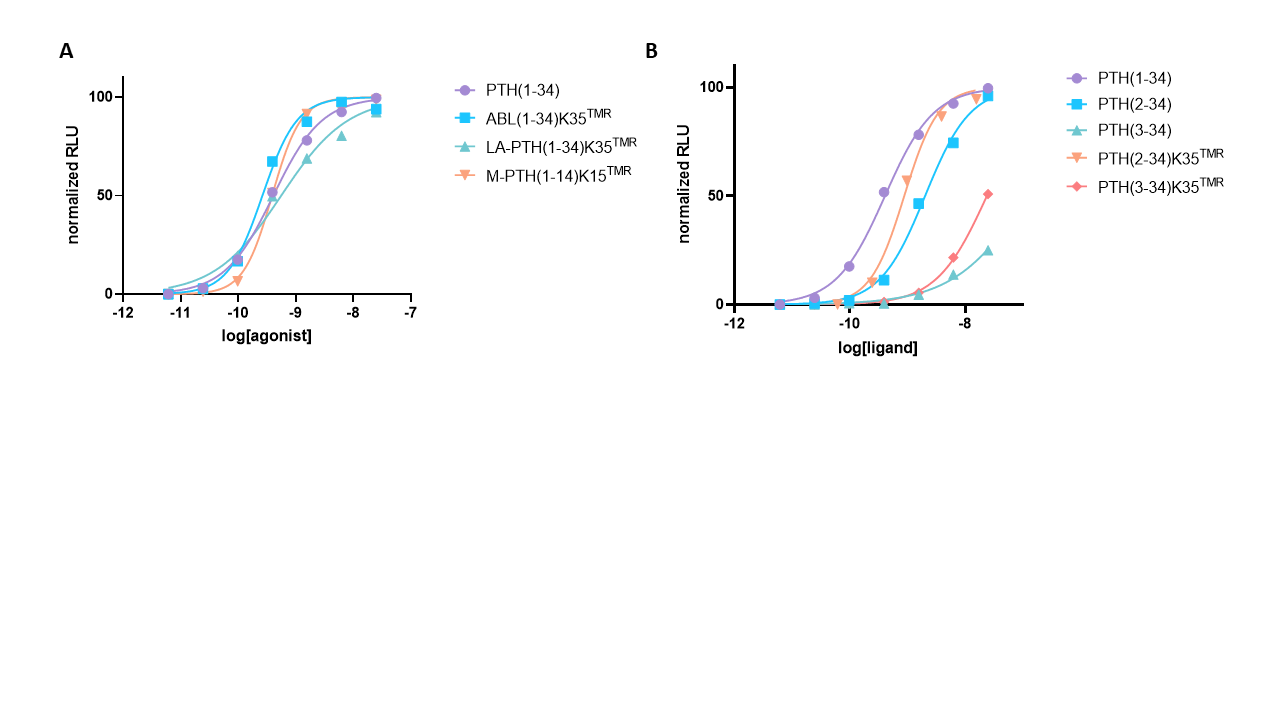

Figure S7. (A) The cAMP formation stimulated by TMR-labeled ABL, LA-PTH and M-PTH analogs were monitored by Glosensor in HEK293 cells. (B) The cAMP formation stimulated by PTH(2-34) and PTH(3-34) with their TMR-labeled analogs were monitored by Glosensor in HEK293 cells. The luminescence response of vehicle control is normalized to 0%, while maximal response of unlabeled PTH(1-34) is normalized to 100%.
